# NCR13 peptide protects soybean against *Cercospora sojina* by multiple modes of action and additive interaction with chemical fungicides

**DOI:** 10.1101/2025.04.29.651315

**Authors:** Ambika Pokhrel, Vishnu Sukumari Nath, James Godwin, Raviraj Kalunke, Meenakshi Tetorya, Kirk J. Czymmek, Dilip M. Shah

**Affiliations:** Donald Danforth Plant Science Center, St Louis, Missouri 63132, USA; Khalifa Center for Genetic Engineering and Biotechnology, United Arab Emirates University, Al Ain, 15551; Advanced Bioimaging Laboratory, Donald Danforth Plant Science Center, St Louis, MO 63132, USA

**Keywords:** *Cercospora sojina*, Frogeye leaf spot disease, Fungicide resistance, Peptide-fungicide synergy, Peptide-based bio-fungicide, Modes of action

## Abstract

Frogeye Leaf Spot (FLS) disease, caused by a fungal pathogen *Cercospora sojina*, is a serious threat to soybean production globally. The control of FLS is facing a major challenge due to the rapid emergence of pathogen resistance to Quinone outside Inhibitor (QoI) fungicides. Effective long-term management of FLS in soybean calls for the discovery of antifungal compounds with new modes of action (MoA), durability and safety. Here, we showed that chickpea nodule-specific cysteine-rich peptide, NCR13_Peptide Folding Variant1 (NCR13_PFV1), exhibited antifungal activity against QoI-sensitive and -resistant field isolates of *C. sojina* at nanomolar concentrations representing the first antifungal NCR peptide reported effective against *C. sojina*. Spray-application of this peptide showed no phytotoxicity and effectively protected soybean against FLS. When combined with the QoI fungicide azoxystrobin, NCR13_PFV1 provided additive control of FLS. NCR13_PFV1 induced plasma membrane disruption and production of reactive oxygen species (ROS) in *C. sojina*. NCR13_PFV1 was rapidly internalized into fungal cells where it accumulated in the cytoplasm, localized inside nucleus, bound to fungal ribosomal RNA and inhibited protein translation *in vivo*. RNA-seq studies revealed the upregulation of several genes encoding heme binding proteins in peptide-challenged *C. sojina*. Notably, iron supplementation in the growth medium reduced the peptide-induced ROS and antifungal activity, revealing the importance of iron homeostasis in protection or recovery of *C. sojina* from oxidative stress. Overall, NCR13_PFV1 with multiple MoA holds potential as a bio-fungicide for FLS control complementing conventional QoI fungicides and overcoming fungicide resistance in *C. sojina*.

## Introduction

Soybean (*Glycine max* Merr.) is a highly versatile economically important oilseed crop grown and consumed worldwide. However, its production is constrained by various fungal diseases (Whitham et al., 2016). Frogeye leaf spot (FLS) caused by *Cercospora sojina* Hara, is one of the most economically important foliar diseases severely affecting soybean production in the United States and other soybean growing areas of the world (Barro et al., 2023). Although FLS has historically been more prevalent in the southern United States, in recent years, its impact on soybean production has increased substantially across the north central states. This geographical expansion of FLS disease has most likely been due to warmer winter temperatures and the ability of *C. sojina* to survive in leaf debris for up to two years (Cruz and Dorrance, 2009; Zhang and Bradley, 2014; Barro et al., 2023).

In the United States, FLS is managed primarily through application of quinone outside inhibitor (QoI, Fungicide Resistance Action Committee [FRAC] group 11) class of fungicides. The QoI fungicides have a single-site mode of action (MoA). They block electron transport to the outer quinol oxidation site of the cytochrome bc1 enzyme complex III, resulting in the interruption of ATP production and inhibition of mitochondrial respiration (Bartlett et al., 2002). The QoI fungicides have provided excellent control of FLS since the commercial introduction of azoxystrobin in 1996; however, their efficacy has since declined rendering them ineffective for control of FLS. The emergence of resistance to QoI fungicides in *C. sojina* is now widespread in several soybean-producing states of the country (Barro et al., 2023). Resistance to QoI fungicides in *C. sojina* is caused by a nonsynonymous substitution of glycine to alanine at position 143 in cytochrome b (Zeng et al., 2015). The widespread resistance in *C. sojina* to QoI fungicides has become a major concern. Therefore, an effective management strategy of FLS calls for the discovery of sustainable bio-fungicides with multiple MoA capable of mitigating fungicide resistance. Small cysteine-rich antifungal peptides (AFP) with multi-faceted MoA hold potential for development as bio-fungicides that could replace or complement the QoI fungicides (Velivelli et al., 2020; Tetorya et al., 2023; Djami-Tchatchou et al., 2023; Li et al., 2024). Understanding the mechanisms underlying the uptake of AFPs and their MoA in targeted fungal pathogens will facilitate their development as bio-fungicides.

The cysteine-rich defensins are the best characterized class of plant AFPs for their antifungal activity and multi-faceted MoA. Many fungal pathogens internalize defensins and defensin-derived peptides *in vitro* making them promising candidates to develop them as bio-fungicides to protect crops. For example, a well-characterized plant defensin MtDef4 and the 17-amino acid γ-core peptide GMA4CG_V6 peptide derived from this defensin are internalized by a necrotroph *Botrytis cinerea in vitro* and robustly reduce gray mold symptoms when sprayed on tomato plants (Li, et al. 2024; Tetorya, et al. 2023). During the last few years, cationic nodule-specific cysteine-rich (NCR) peptides expressed in legumes of the inverted-repeat-lacking-clade (IRLC) have emerged as a new class of AFPs with potent antifungal activity against plant and human fungal pathogens (Ördögh et al., 2014; Velivelli et al., 2020; Lima et al., 2022). NCR peptides are predicted to form 2 or 3 disulfide bonds (Maróti et al., 2015; Mergaert, 2018). The chickpea NCR13 is a 32-amino acid peptide predicted to form three disulfide bonds (Godwin et al., 2024). NCR13 expressed in *Pichia pastoris* formed two disulfide cross-linking variants, NCR13_Peptide Folding Variant1 (NCR13_PFV1) and NCR13_PFV2, that had an identical amino acid sequence but differed in the arrangement of the two out of three disulfide bonds.

NCR13_PFV1 exhibited several fold more potent antifungal activity than NCR13_PFV2 against *B. cinerea, Fusarium virguliforme,* and *Sclerotinia sclerotiorum*. These two peptides also displayed strikingly different MoA against *B. cinerea* (Godwin et al., 2024).

No AFP with inhibitory activity against *C. sojina* has been reported to date. In this study, we tested several defensin-derived and NCR AFPs and found that NCR13_PFV1 was most potent against both QoI-sensitive and -resistant field isolates of *C. sojina* with a minimal inhibitory concentration (MIC) at nanomolar concentrations. Spray-application of NCR13_PFV1 on the leaves of soybean plants greatly reduced FLS disease symptoms. Furthermore, peptide-fungicide combination assays revealed that NCR13_PFV1 exhibited additive antifungal activity in combination with the QoI fungicide azoxystrobin against *C. sojina in planta* providing robust control of FLS. NCR13_PFV1 exhibited multiple MoA against *C. sojina* and was non-phytotoxic to soybean plants. The RNA-seq study revealed the importance of iron homeostasis in alleviating peptide-induced oxidative stress and protecting fungus from this peptide. Our findings suggest that NCR13_PFV1 could be a powerful tool for the sustainable management of FLS in soybean fields.

## Results

### NCR13_PFV1 exhibits potent antifungal activity against the QoI fungicide-sensitive and - resistant *C. sojina* isolates

To identify AFPs highly active against QoI fungicide-sensitive and -resistant field isolates of *C. sojina*, we screened NCR13_PFV1, NCR13_PFV2, and NCR13-derived short peptide variants NCR13_V7 and NCR13_V9 (Fig. 1a). Several variants of the short defensin-derived peptides (Fig. 1b) were also tested during the initial screen. The MICs of defensin-derived peptides ranged from 3 to 12 µM (Fig. 1b). Importantly, NCR13_PFV1 showed the most potent inhibitory activity against multiple fungicide-sensitive and -resistant isolates of *C. sojina* with MIC of 0.18 µM (Fig. 1a). Notably, NCR13_PFV2 was four- to eight-fold less potent with the MIC of 0.75-1.5 µM (Fig. 1a). Due to NCR13_PFV1’s potent antifungal activity against *C. sojina*, we focused on characterizing MoA and *in planta* antifungal activity of NCR13_PFV1 against the QoI fungicide-resistant *C. sojina* either alone or in combination with the fungicides.

**Figure 1:**
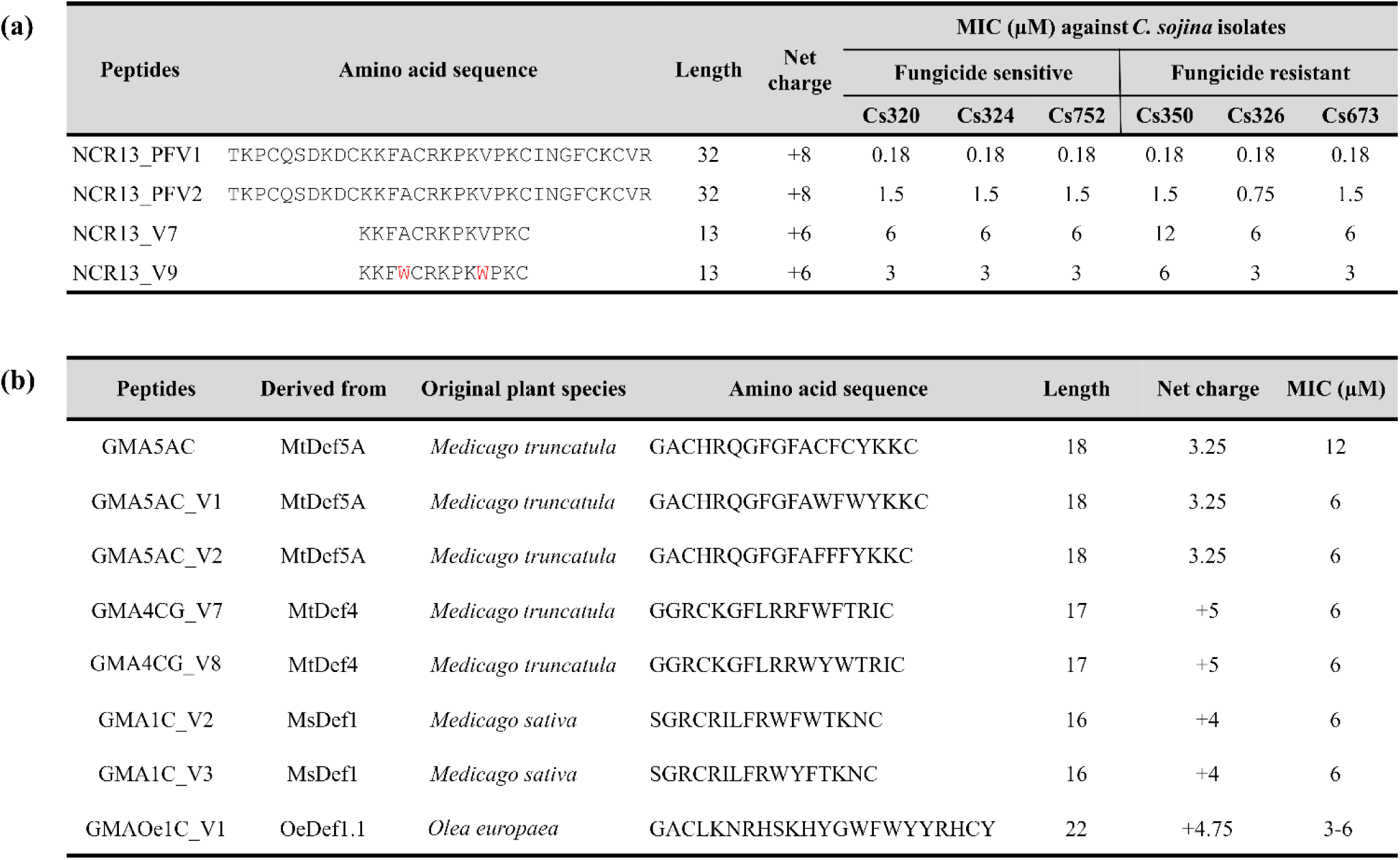
NCR13 and defensin derived peptides exhibit *in vitro* antifungal activity against *C. sojina* quinone outside inhibitor (QoI) fungicide-sensitive and -resistant isolates. (a) Amino acid sequence, length, net charge and *in vitro* MIC of NCR13 and its variants against various QoI fungicide sensitive and resistant field isolates of *C. sojina*. **(b)** Amino acid sequence, length, net charge and MIC of various defensin derived peptides against QoI fungicide resistant isolate Cs350 of *C. sojina*. The MIC of each peptide was determined using both resazurin cell viability assay and microscopy.

### Pre- and post-inoculation spray-application of NCR13_PFV1 on soybean plants reduces FLS disease symptoms

To test the ability of spray-applied NCR13_PFV1 (6 µM) to reduce FLS disease symptoms *in planta*, the peptide was spray-applied on the leaves of two- to three-week-old old soybean plants challenged with the QoI fungicide-sensitive and -resistant *C. sojina* isolates Cs320 and Cs350, respectively. For pre-inoculation *in planta* antifungal activity assay, soybean leaves were sprayed with 1 mL of 6 µM NCR13_PFV1 followed by 1 mL of Cs320 or Cs350 conidia 2 hr later. For post-inoculation *in planta* antifungal assay, soybean leaves were first inoculated with 1 mL of fungal conidia followed by spraying with 6 µM NCR13_PFV1 2 hr later. NCR13_PFV1 applied pre- or post-inoculation on soybean leaves *in planta* significantly reduced (*p*-value ≤ 0.05) FLS symptoms caused by fungicide-sensitive Cs320 or -resistant Cs350 isolates (Fig. 2a, b, d, e).

**Figure 2:**
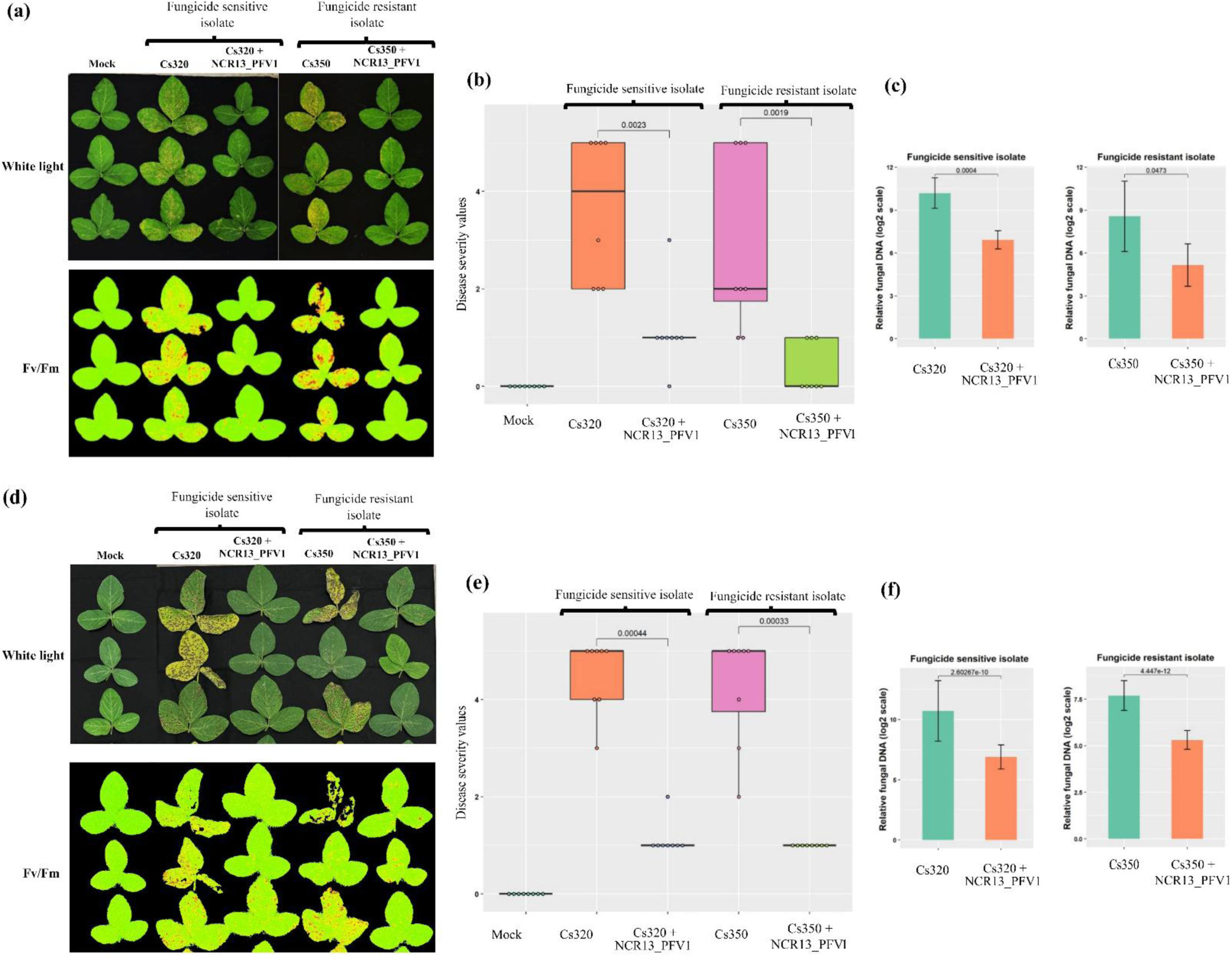
NCR13_PFV1 applied pre- and post-inoculation reduces FLS disease symptoms in soybean. **(a,d)** White light and CropReporter (Fv/Fm) images of *in planta* (**a**) pre-inoculation and (**d**) post-inoculation antifungal activity of 6 μM NCR13_PFV1 against *C. sojina*. The inoculated leaves of each plant were detached and imaged with normal light as well as on CropReporter. Red color in CropReporter image represents extensive tissue damage and low photosynthetic efficiency (Fv/Fm) whereas green color indicates healthy tissue and higher photosynthetic efficiency (N=8 and R=2, where N refers to number of plants and R refers to number of biological replicates). **(b, e)** Box plots showing the disease severity values of FLS disease on soybean leaves from (**b**) preinoculation and (**e**) post-inoculation assays. The disease rating was given based on a 0-5 scale, where 0 was a healthy plant with no FLS symptoms and 5 indicated a severely diseased or dead leaf. **(c, f)** The bar graphs showing the relative fungal DNA content in the Cs320 and Cs350 *C. sojina* treated soybean leaves in the presence and absence of NCR13_PFV1 as determined by qPCR from both (**c**) pre-inoculation and (**f**) post-inoculation *in planta* assays. Data are shown as the mean of two biological replicates, each with 4 plants and error bars indicating standard error of the mean between the replicates. The statistical test was performed using the Wilcoxon test and the *p*-values among the compared groups are indicated. *p*-value ≤ 0.05 is considered as statistically significant.

Furthermore, both Cs320 and Cs350 isolates had significantly reduced biomass (*p*-value ≤ 0.05) in the infected soybean leaves sprayed with the peptide as revealed by fungal DNA quantification using qPCR. (Fig. 2c, f). Overall, since NCR13_PFV1 foliar spray controls FLS symptoms in soybean leaves, it has potential to be developed as bio-fungicide.

### NCR13_PFV1 and QoI fungicide combinations provide additive antifungal activity against *C. sojina in vitro*

Since *C. sojina* has already developed resistance to QoI fungicides (Zeng et al., 2015), we aimed to evaluate whether combining NCR13_PFV1 with existing QoI fungicides could rescue their activity against *C. sojina*. To determine the efficacy of combinations between NCR13_PFV1 and three QoI fungicides, azoxystrobin, trifloxystrobin, and pyraclostrobin against the fungicide-sensitive isolate Cs320 and the fungicide-resistant isolate Cs350, checkerboard assays were performed. The MIC values of trifloxystrobin and pyraclostrobin against fungicide sensitive isolate Cs320 was extremely low (<0.02 µM), where 0.02 µM was the lowest concentration tested (Table S1). Therefore, checkerboard assays to determine antifungal activity of NCR13_PFV1 in combination with trifloxystrobin or pyraclostrobin against Cs320 were not performed. The MIC values of the peptide, fungicide, or peptide plus fungicide were recorded after 96 hr (4 days). NCR13_PFV1 and all tested QoI fungicides (azoxystrobin, trifloxystrobin, and pyraclostrobin) exhibited additive antifungal activity against *C. sojina* Cs350 *in vitro* with the fractional inhibitory concentration (FIC) score of 1 or less (Fig. 3a). Similarly, NCR13_PFV1 in combination with azoxystrobin also exhibited additive antifungal activity against fungicide-sensitive *C. sojina* Cs320 (Fig. 3a). When combined, the MIC of fungicides as well as NCR13_PFV1 was reduced two-fold or more. Since the highest concentration of fungicide tested was 48 µM, the actual MIC values against *C. sojina* were noted >48 µM (Fig. 3a, S1). Overall, checkerboard assays clearly showed that NCR13_PFV1 and QoI fungicides demonstrated additive antifungal activity against both fungicide-sensitive and -resistant isolates *in vitro*.

**Figure 3:**
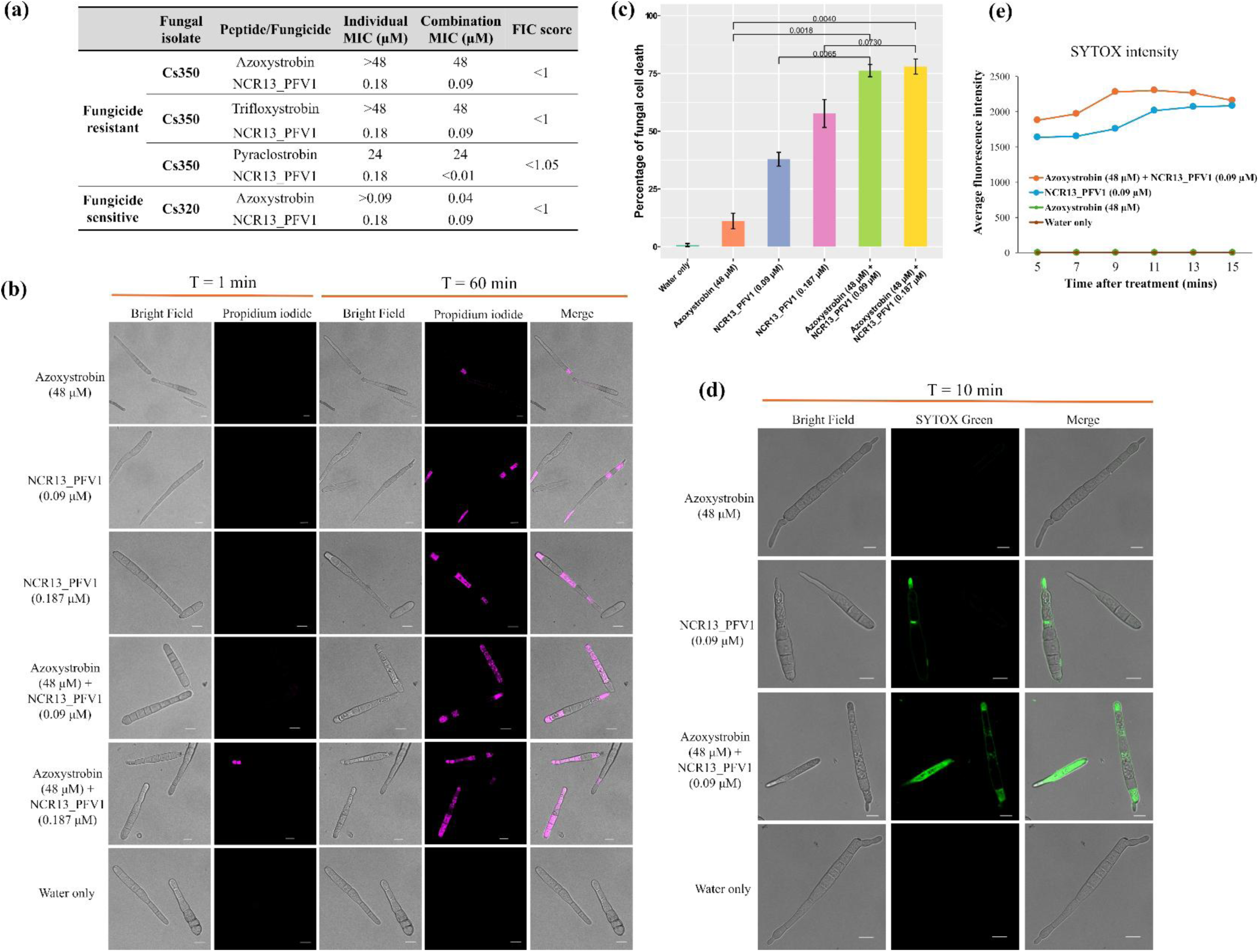
NCR13_PFV1 and quinone outside inhibitor (QoI) fungicide combinations additively increase their antifungal activity against *C. sojina in vitro*. (a) The individual as well as combined MIC of NCR13_PFV1 and various QoI fungicides against *C. sojina* isolates in checkerboard assays as determined in 4 days. MIC was determined using microscopy. FIC indicates the fraction inhibitory concentration. The FIC index was interpreted as Synergistic: ≤ 0.5; Additive: > 0.5-1.0; Indifferent: >1-< 4.0; Antagonistic: ≥ 4.0. **(b**) Representative time-lapse confocal microscopy images of *C. sojina* Cs350 conidia showing cell death at 1 and 60 min after treatment of azoxystrobin, NCR13_PFV1, and their combinations. Scale bars = 10 μm. **(c)** Bar graph showing the percentage of cell death in *C. sojina* Cs350 determined at 1 hr after challenge with various combinations of peptide and fungicide. Cell death indicator dye propidium iodide (PI) was used to monitor cell death in Cs350. The statistical test was performed using the Wilcoxon test and *p*values among the compared groups are indicated. *p*-value ≤ 0.05 is considered statistically significant. **(d)** Time-lapse confocal microscopy images of SYTOX Green (SG) uptake in Cs350 treated with azoxystrobin, NCR13_PFV1, and their combinations at 10 min. Scale bars = 10 μm. **(e)** Quantitative measurements of average fluorescence intensity of SYTOX Green over time to assess membrane permeabilization in various combinations of peptide and fungicide in *C. sojina* Cs350.

Among the QoI fungicides, we focused on azoxystrobin because it is the most widely used fungicide. The combination of azoxystrobin and NCR13_PFV1 reduced the MIC of both at least two-fold against *C. sojina*. We hypothesized that this additive antifungal activity could be due to rapid fungal cell death that could happen very early on in the combination treatments. To further explore the effect of azoxystrobin and NCR13_PFV1 combination in causing cell death in fungicide resistant isolate Cs350 at early timepoints (within 1 hr), propidium iodide (PI) staining was used. We observed that Cs350 independently treated with azoxystrobin (48 μM) and NCR13_PFV1 (0.18 μM) resulted in 11% and 57% of the cell death within 1 hr, respectively (Fig. 3b, c). However, the combination of azoxystrobin (48 μM) and NCR13_PFV1 (0.18 μM) increased the cell death to 78% in 1 hr. Similarly, azoxystrobin (48 μM) and NCR13_PFV1 (0.09 μM) combination also increased the cell death to 76% at 1 hr. It is important to point out that the combination MIC of azoxystrobin and NCR13_PFV1 against *C. sojina* Cs350 at 4 days was 48 μM and 0.09 μM, respectively (Fig. 3a). Overall, combination of NCR13_PFV1 and azoxystrobin significantly promotes cell death in *C. sojina* Cs350 at 1 hr after treatment, suggesting that additive antifungal activity of NCR13_PFV1 and azoxystrobin is due to rapid cell death.

The checkerboard and cell death assays confirmed the additive antifungal activity of azoxystrobin and NCR13_PFV1 against *C. sojina;* however, how this peptide and fungicide complement each other needs to be further explored. One of the important MoA of NCR13_PFV1 is permeabilization of plasma membrane in *C. sojina* conidia and mycelia (Fig. S2). Therefore, we hypothesized that NCR13_PFV1 permeabilizes the plasma membrane in *C. sojina* facilitating QoI fungicides including azoxystrobin to enter inside the fungal cells much faster and cause quicker cell death. Using the SYTOX Green (SG) plasma membrane permeabilization assay, we found that azoxystrobin alone did not permeabilize the plasma membrane. However, the fungicide in combination with NCR13_PFV1 caused greater membrane permeabilization, as indicated by higher SG fluorescence intensity compared to the peptide alone (Fig. 3d, e). This further confirmed the additive antifungal activity of azoxystrobin and NCR13_PFV1 combination against *C. sojina* Cs350 and opens a new possibility of crop protection against FLS through foliar application of peptide-fungicide combination. Due to the importance of fungicide resistance in *C. sojina*, all experiments described hereafter were conducted using the fungicide-resistant isolate Cs350.

### Additive reduction of FLS disease severity in soybean by NCR13_PFV1 and azoxystrobin combination

To further assess additive antifungal activity between NCR13_PFV1 and azoxystrobin, post-inoculation *in planta* antifungal assays were performed at various concentrations of both peptide and fungicide. These assays showed that compared to the individual spray-application of NCR13_PFV1 or azoxystrobin on soybean leaves, spray-application of the peptide-fungicide combination reduced the FLS symptoms to a significantly higher level (Fig. 4). Even though the MIC of azoxystrobin *in vitro* was >48 µM (Fig. 3a), it was still able to reduce FLS symptoms *in planta* when sprayed at concentrations of 3 and 6 µM (Fig. 4a). Azoxystrobin at low concentrations of 3 and 6 µM does not inhibit fungal growth *in vitro*, but it may be able to reduce disease severity by preventing conidial germination or reducing hyphal growth *in planta* in concert with plant defense mechanisms. Similarly, peptide-fungicide combination at 3 µM NCR13_PFV1 and 6 µM azoxystrobin provided significantly better control of FLS than the combination at 3 µM each (Fig. 4). Based on these findings, we conclude that NCR13_PFV1 and azoxystrobin combinations provide more robust control of FLS *in planta* than the peptide or fungicide alone.

**Figure 4:**
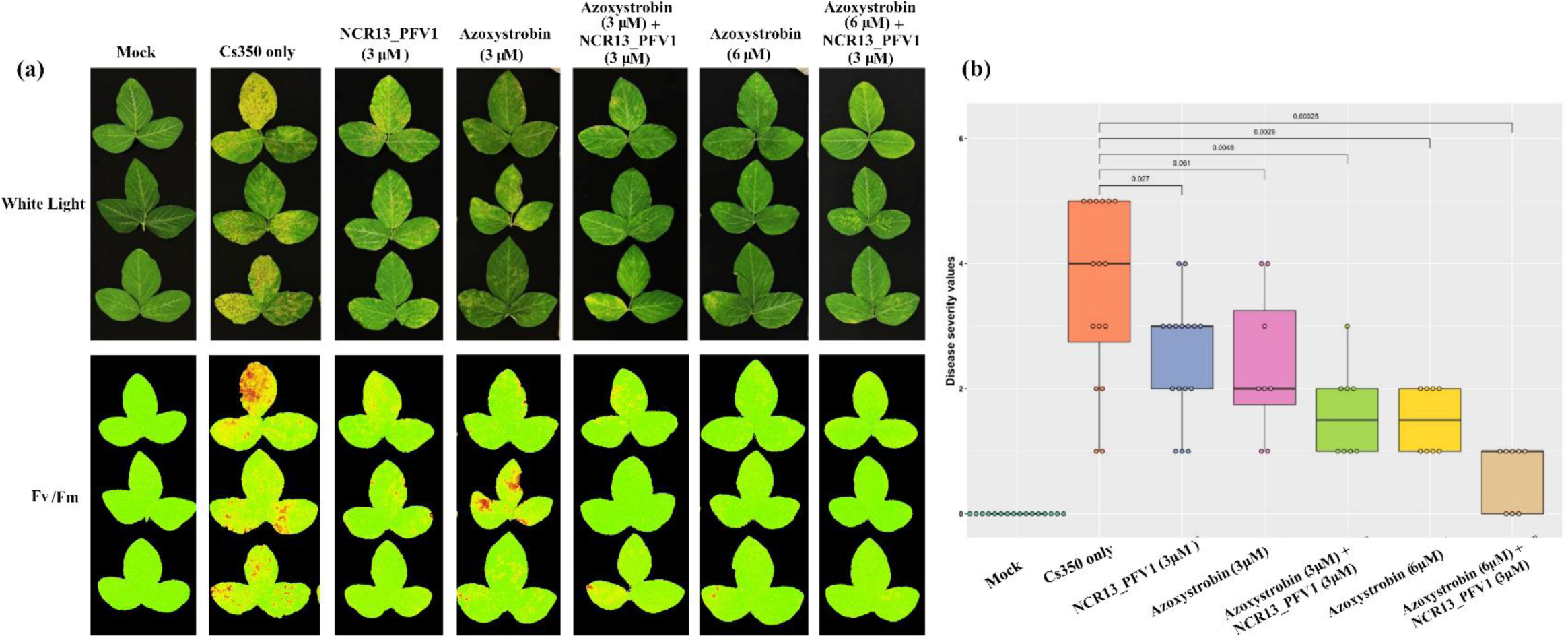
NCR13_PFV1 and azoxystrobin combinations provide robust control of FLS disease in soybean. **(a)** White light and CropReporter (Fv/Fm) images of *in planta* post-inoculation antifungal activity of NCR13_PFV1 and azoxystrobin against Cs350 *C. sojina*. The inoculated leaves of each plant were detached and imaged with normal light as well as on CropReporter. Red color in CropReporter image represents extensive tissue damage and low photosynthetic efficiency (Fv/Fm) whereas green color indicates healthy tissue and higher photosynthetic efficiency (N=8-16 and R=2, where N refers to number of plants and R refers to number of biological replicates). **(b)** The box plots showing the disease severity values of FLS disease on soybean leaves at various concentrations of NCR13_PFV1 and azoxystrobin. The disease rating was given based on a 0-5 scale, where 0 was a healthy plant with no FLS symptoms and 5 indicated a severely diseased or dead leaf. The Statistical test for significance was performed using the Wilcoxon test and the *p*-values among the compared groups are indicated. *p*-value ≤ 0.05 is considered statistically significant.

### NCR13_PFV1 is localized in cytoplasm and nucleus in *C. sojina*

To determine the subcellular localization of NCR13_PFV1 in *C. sojina*, confocal microscopy was done using DyLight550-labeled NCR13_PFV1. We found that DyLight550-labeled NCR13_PFV1 was internalized by *C. sojina* conidia within 1 min and accumulated in the cytoplasm (Fig. 5a, S3, S4). Additionally, NCR13_PFV1 localized into the nucleus (Fig. 5b, white arrows) and other intensely stained regions in *C. sojina*. Notably, this nuclear localization was mostly observed at 0.09 μM (0.5 x MIC) of NCR13_PFV1. Further studies to identify the intensely stained regions revealed no evidence of co-localization of the peptide with either lipid bodies or vacuole (Fig. S3, S4).

**Figure 5:**
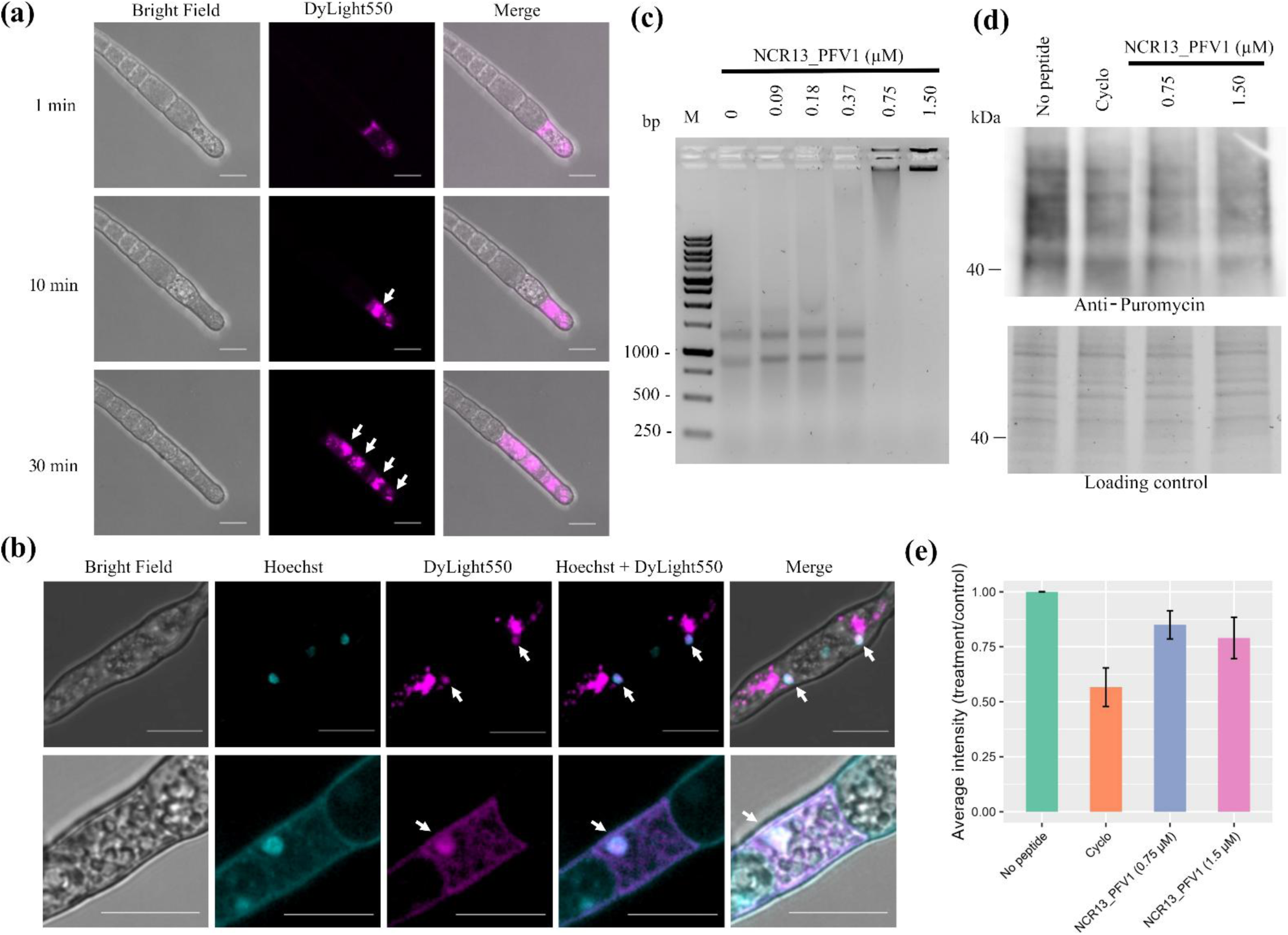
NCR13_PFV1 is internalized in *C. sojina*, localizes in cytoplasm and nucleus, binds to the rRNA, and inhibits protein translation. **(a)** Time-lapse confocal microscopy images of *C. sojina* showing the internalization of 0.187 μM of NCR13_PFV1 after 1, 10, and 30 min of treatment. The white arrow indicates the aggregation of peptide in the cytoplasm. Scale bars = 10 μm. **(b)** Confocal microscopy images of *C. sojina* showing internalization of 0.09 μM of NCR13_PFV1 after 30 min of treatment. The white arrow indicates NCR13_PFV1 colocalization inside nucleus. Scale bars = 10 μm). **(c)** rRNA binding activity of NCR13_PFV1 tested using gel electrophoresis mobility shift assay (EMSA). The *C. sojina* RNA (∼200 ng) was incubated with different concentrations of NCR13_PFV1 and visualized in 1% agarose gel. The first lane on the gel M is a 1Kb DNA ladder. **(d)** Protein translation inhibition assessed using puromycin labeling assay. Protein translation inhibition and total protein content in *C. sojina* germlings in presence of water (no peptide), cycloheximide (Cyclo), and different concentrations of NCR13_PFV1. Upper panel shows the puromycin-labeled nascent polypeptides in *C. sojina* germlings detected by anti-puromycin antibody and the bottom panel shows the stain-free gel image showing equal total protein loading across all lanes. **(e)** The bar graph shows the average intensity of protein translation inhibition calculated for each treatment (n=3).

### NCR13_PFV1 binds to *C. sojina* ribosomal RNA and inhibits protein translation *in vivo*

In our previous study we demonstrated that NCR13_PFV1 binds to rRNA and inhibits protein translation in *B. cinerea* germlings (Godwin et al., 2024; Godwin and Shah, 2025). Therefore, we hypothesized that NCR13_PFV1 might have similar MoA in *C. sojina.* Electrophoretic mobility shift assay (EMSA) revealed that NCR13_PFV1 binds to *C. sojina* rRNA (Fig. 5c) consistent with its binding ability with *B. cinerea* rRNA (Godwin et al., 2024). While the MIC of NCR13_PFV1 against *C. sojina* is 0.187 μM, the binding of NCR13_PFV1 with rRNA was observed starting at 0.75 μM. Additionally, using a puromycin labelling assay (Godwin and Shah, 2025), we found that NCR13_PFV1 inhibited protein translation *in vivo* in *C. sojina,* as shown by reduced levels of puromycylated peptides, which reflect an overall decrease of protein translation (Fig. 5d,e). While the MIC of this peptide against *C. sojina* is 0.187 μM, protein translation inhibition in *C. sojina* germlings was observed at higher concentrations, starting at 0.75 and 1.5 μM. These results suggest that binding with rRNA and inhibiting protein translation could be an important MoA of NCR13_PFV1 against multiple fungal pathogens.

### NCR13_PFV1 treatment induces the expression of genes encoding heme binding proteins in *C. sojina*

To gain further insight into the MoA of NCR13_PFV1, we performed RNA-seq study on *C. sojina* germlings challenged with the 0.187 μM peptide. Among 1,142 differentially expressed genes (DEGs) identified after 30 min of peptide treatment in *C. sojina,* 645 genes were upregulated, and 497 genes were downregulated (Fig. 6a; Table S2, S3). The significantly upregulated genes include several classes of transporters, peroxidases, dehydrogenases, transcription factors, proteins of unknown function (Table S2). Interestingly, a gene encoding the respiratory burst oxidase homolog (RBOH) which is known to be crucial for production of ROS, was among the top five significantly upregulated genes (Fig. 6b). The gene ontology (GO) enrichment analyses of DEGs revealed several GO terms enriched in *C. sojina* upon peptide treatment (Fig. 6c, d). Among enriched GO terms, heme binding and tetrapyrrole binding were significantly upregulated (Fig. 6c; Table S4). Both tetrapyrrole binding and heme binding contained the same list of 16 different genes encoding cytochrome P450s, catalases, peroxidase, etc. (Fig. 6d). Among downregulated genes 15 GO terms related to biological process and 5 terms related to metabolic process were significantly enriched (Fig. 6e; Table S5). In comparison with the NCR13_PFV1 treatment, we found that NCR13_PFV2 treatment also induced genes related to heme and tetrapyrrole binding in *C. sojina* (Fig. S5). Strikingly, similar results were observed in a different fungal pathogen, *B. cinerea,* in response to NCR13_PFV1 treatment using transcriptomic analyses (Fig. S6). Overall, our findings suggest that the upregulation of genes encoding heme binding proteins could be a conserved mechanism adopted by various fungi to protect against NCR13 as well as other peptides.

**Figure 6:**
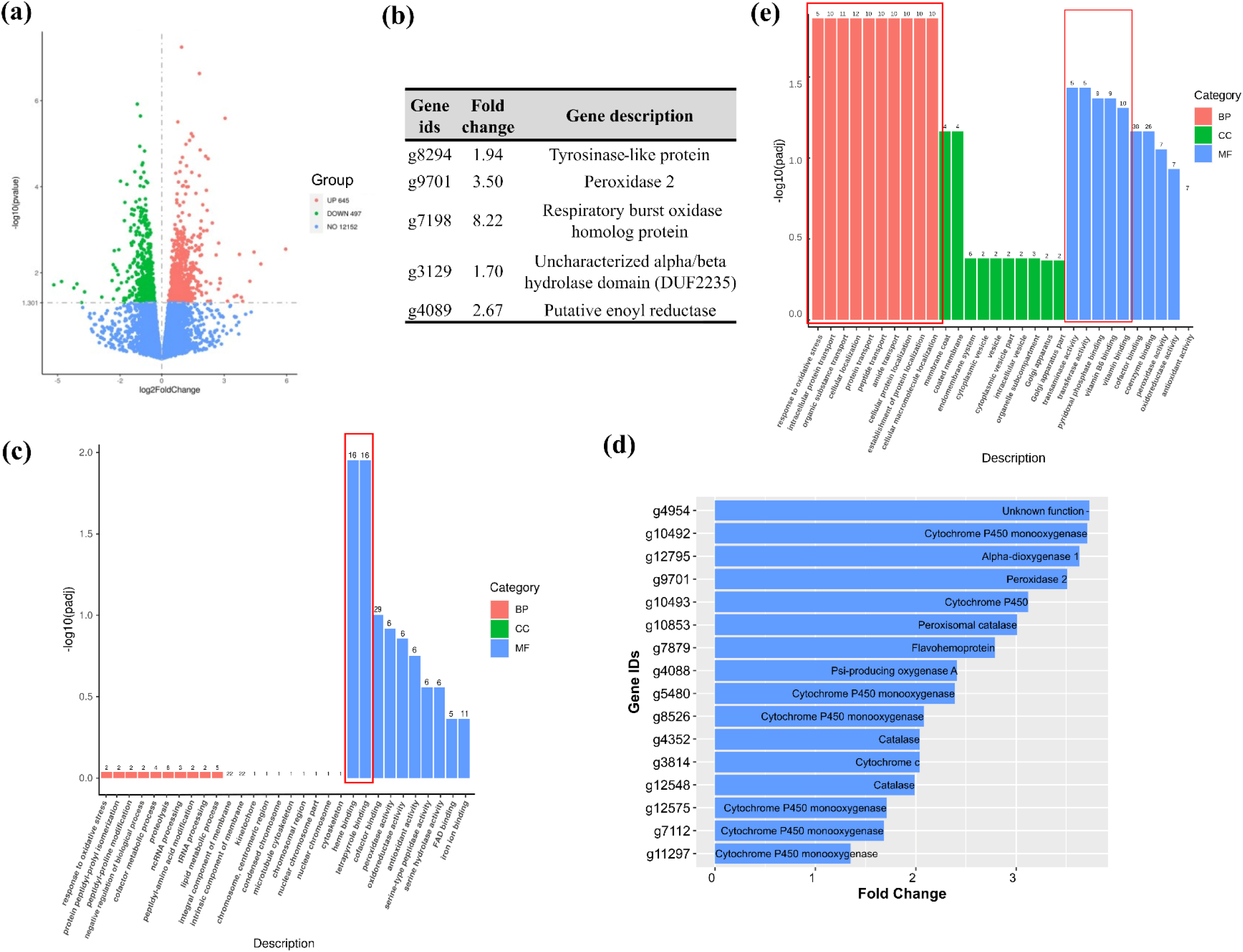
NCR13_PFV1 upregulates genes encoding heme binding proteins in *C. sojina*. **(a)** Volcano plot showing the number of differentially expressed genes in *C. sojina* after 30 min of NCR13_PFV1 treatment when compared to the no peptide control at 0 min. **(b)** Table showing the top five significantly upregulated genes (sorted based on *p*-adjusted values) in *C. sojina* after NCR13_PFV1 treatment. A gene potentially involved in the induction of ROS in *C. sojina* is highlighted in red. **(c, e)** Bar graphs showing the top ten **(c)** upregulated and **(e)** downregulated GO terms enriched in each category (BP-Biological Process, CC-Cellular Component, and MF-Molecular Function) at 30 min after NCR13_PFV1 treatment in *C. sojina*. The significantly enriched GO terms at *p*-value≤0.05 are highlighted in the red rectangular box. **(d)** List of 16 upregulated genes involved in the heme and tetrapyrrole binding in *C. sojina* after NCR13_PFV1 treatment.

### NCR13 does not bind with heme

RNA-seq data showed upregulation of genes encoding heme binding proteins in *C. sojina* following NCR13_PFV1 treatment and some NCRs, such as NCR247 and NCR455, were previously shown to bind with heme (Sankari et al., 2022). Therefore, we hypothesized that NCR13_PFV1 could bind with heme causing iron deficiency and upregulation of genes encoding heme binding proteins. To assess heme binding ability of NCR13_PFV1 and NCR13_PFV2, we used the previously characterized heme binding peptides, NCR247 and NCR455, as positive controls and performed the heme binding assay (Sankari et al., 2022. A visible color change of heme from brown to reddish-brown was observed upon treatment with NCR247 and NCR455 (Fig. 7a), confirming heme binding. However, no such color change was observed when heme was incubated with NCR13_PFV1 or NCR13_PFV2 (Fig. 7a), suggesting that NCR13 folding variants do not bind heme under the tested conditions. To further verify the lack of heme binding ability of NCR13_PFV1, we performed peroxidase assay. Upon binding to NCR247, heme becomes chemically inaccessible for peroxidase activity. We found that the peroxidase activity of heme and FeSO4, in the presence of NCR13_PFV1 or NCR13_PFV2 was not reduced or stopped as in NCR247 and NCR455 treated positive control samples (Fig. 7b, S7a). These results suggested that NCR13 is not a heme binding peptide.

**Figure 7:**
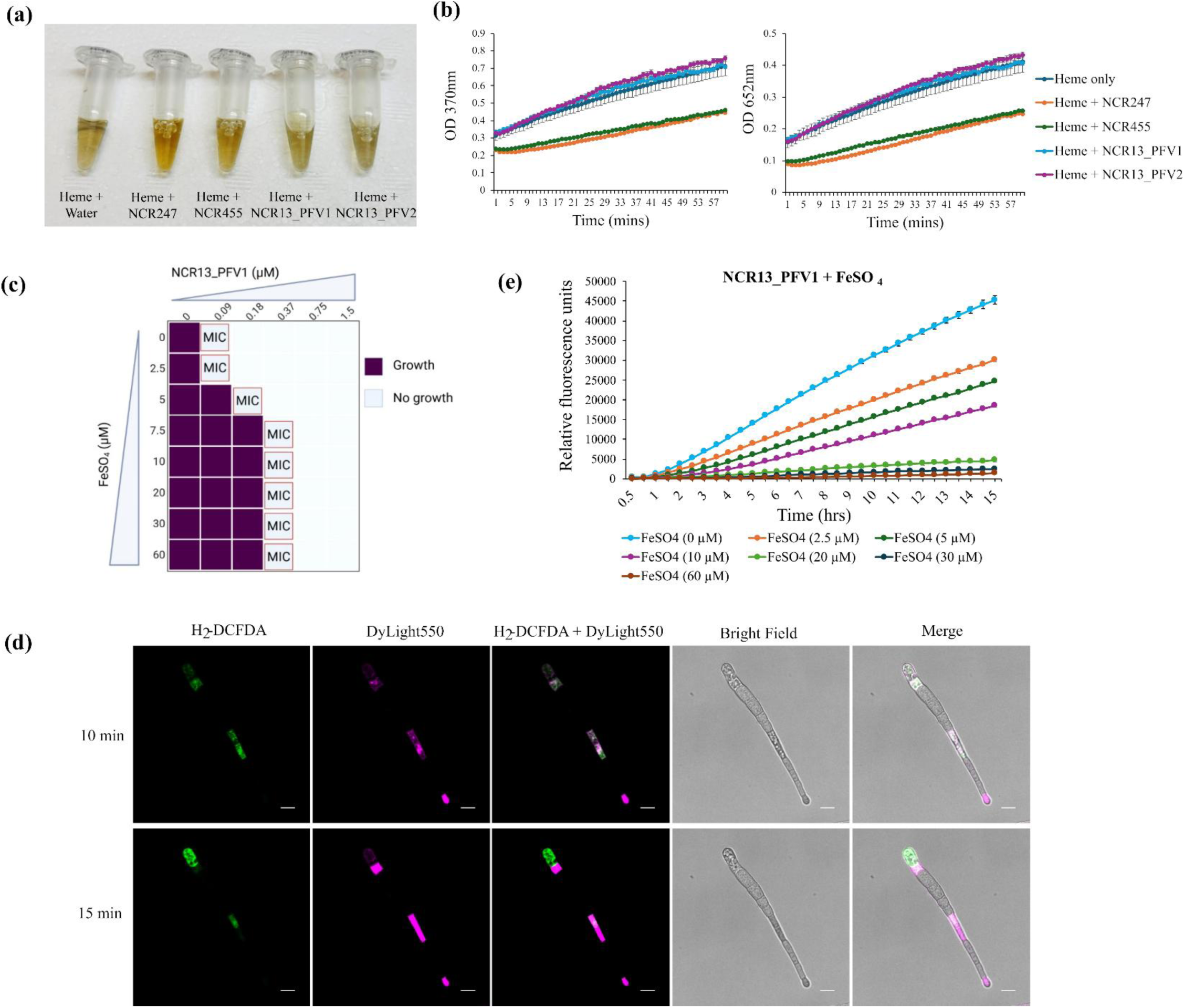
*C. sojina* exhibits iron mediated tolerance to NCR13_PFV1. **(a)** NCR247 and NCR455 turn reddish brown following heme binding, whereas NCR13_PFV1, NCR13_PFV2, and heme only treatment shows no color change. **(b)** Graphs show the peroxidase activity of heme and peptides, NCR247, NCR455, NCR13_PFV1, and NCR13_PFV2 on a chromogenic substrate, 3,3′,5,5′-tetramethylbenzidine (TMB), measured by absorption at 370 nm and 652 nm. **(c)** Plot showing depiction of *in vitro* antifungal activity of NCR13_PFV1 against *C. sojina* at various increasing concentrations of iron (FeSO4) determined at two days after incubation. **(d)** Time-lapse confocal microscopy images of *C. sojina* showed internalization of 0.187 μM NCR13_PFV1 and induction of ROS after 10 and 15 min of treatment. Scale bars = 10 μm. **(e)** Graph with the average relative fluorescence intensity of ROS in *C. sojina* after 0.187 μM of NCR13_PFV1 treatment in various iron supplemented media, measured using Tecan plate reader every 30 min for 15 hr. The ROS indicator dye H2DCFDA at 15 μM was used.

### Iron supplementation alleviates NCR13_PFV1 induced oxidative stress in *C. sojina*

Since NCR13_PFV1 induces the expression of genes encoding heme binding proteins, we hypothesized that *C. sojina* might sequester more iron to protect and recover from oxidative stress or damage caused by the peptide treatment. To evaluate this hypothesis, we performed *in vitro* antifungal activity assays against *C. sojina* in synthetic fungal medium (SFM) with concentrations of iron (FeSO4) ranging from 0 to 60 μM. Notably, the concentration of iron in standard SFM medium used for *in vitro* antifungal activity assays is 5 μM. We found that the MIC of NCR13_PFV1 against *C. sojina* increased from 0.09 μM in SFM without iron to 0.375 μM in SFM with ≥7.5 μM iron (Fig. 7c, S7b). We observed a similar effect with various NCR13 variants (NCR13_PFV2, NCR13_V7, and NCR13_V9), where the MIC of peptide increased two-fold with increasing iron concentration (Fig. S7c). Since the MIC of NCR13_PFV1 against *C. sojina* increased with increasing concentrations of iron, we conclude that reduced iron levels made this fungus more sensitive to peptides, while increased iron made it more tolerant. To determine whether this effect was specific to iron, we conducted similar assays using various concentrations of zinc (ZnSO4). No change in MIC was observed with zinc supplementation, (Fig. S7d), suggesting that the recovery or protection of *C. sojina* from NCR13_PFV1 induced stress is iron specific. In *B. cinerea* treated with NCR13_PFV1 in medium containing increasing concentration of iron from 5 to 30 μM, we observed a similar four-fold increase in MIC (from 0.09 to 0.375 μM) with increasing concentrations of iron (Fig. S8). Based on these results, we conclude that iron plays an important role in providing recovery or protection from AFP-imposed stress in *C. sojina* and *B. cinerea*.

Since ROS is induced upon cellular stress, and our transcriptomic analysis revealed upregulation of several genes encoding heme binding proteins associated with ROS detoxification (Pradhan et al., 2017; Zámocký et al., 2020; Park et al., 2023; Wang et al., 2024), we examined whether NCR13_PFV1 treatment induces ROS in *C. sojina*. The *C. sojina* conidia challenged with DyLight550-NCR13_PFV1 was used in combination with the ROS-detection dye, H₂DCFDA to detect ROS. The ROS accumulation started immediately after internalization of the peptide and the signal intensity peaked right before cell death or cell became saturated with the peptide (Fig. 7d).

To further monitor ROS dynamics in *C. sojina* conidia challenged with 0.187 μM of NCR13_PFV1 in medium containing varying levels of iron, H₂DCFDA fluorescence measurements were taken every 30 min for 15 hr. The ROS levels in peptide-treated conidia were dependent on iron concentration (Fig. 7e). Increasing concentrations of iron reduced the ROS levels in peptide-challenged *C. sojina*. Specifically, at concentrations of iron above 20 μM, the amount of ROS didn’t increase with time (Fig. 7e). Overall, *C. sojina* treated with NCR13_PFV1 accumulates ROS, and elevated level of iron in the growth medium reduces peptide-induced oxidative stress.

### NCR13_PFV1 foliar application does not affect soybean growth and yield

To test if foliar application of NCR13_PFV1 has any negative effects on growth and yield of soybean, three-month-old plants were sprayed with NCR13_PFV1 (bi-weekly from 4^th^ to 10^th^ week) in the greenhouse. They were examined morphologically for defects in aerial plant growth and changes in leaf color. No morphological difference was observed between water treated and NCR13_PFV1 treated plants for parameters such as growth rate, height, and changes in leaf color (Fig. 8a). We found no significant differences in the number of pods, 100 seed weight as well as total seed yield between water and NCR13_PFV1-sprayed plants at maturity as determined by Student’s *t* test (Fig. 8b). Additionally, DyLight550-NCR13_PFV1 sprayed on soybean leaves did not enter inside the leaf epidermal cell cytoplasm but remained on the surface (Fig. 8c). Overall, NCR13_PFV1 treatment had no deleterious effects on soybean growth and yield.

**Figure 8:**
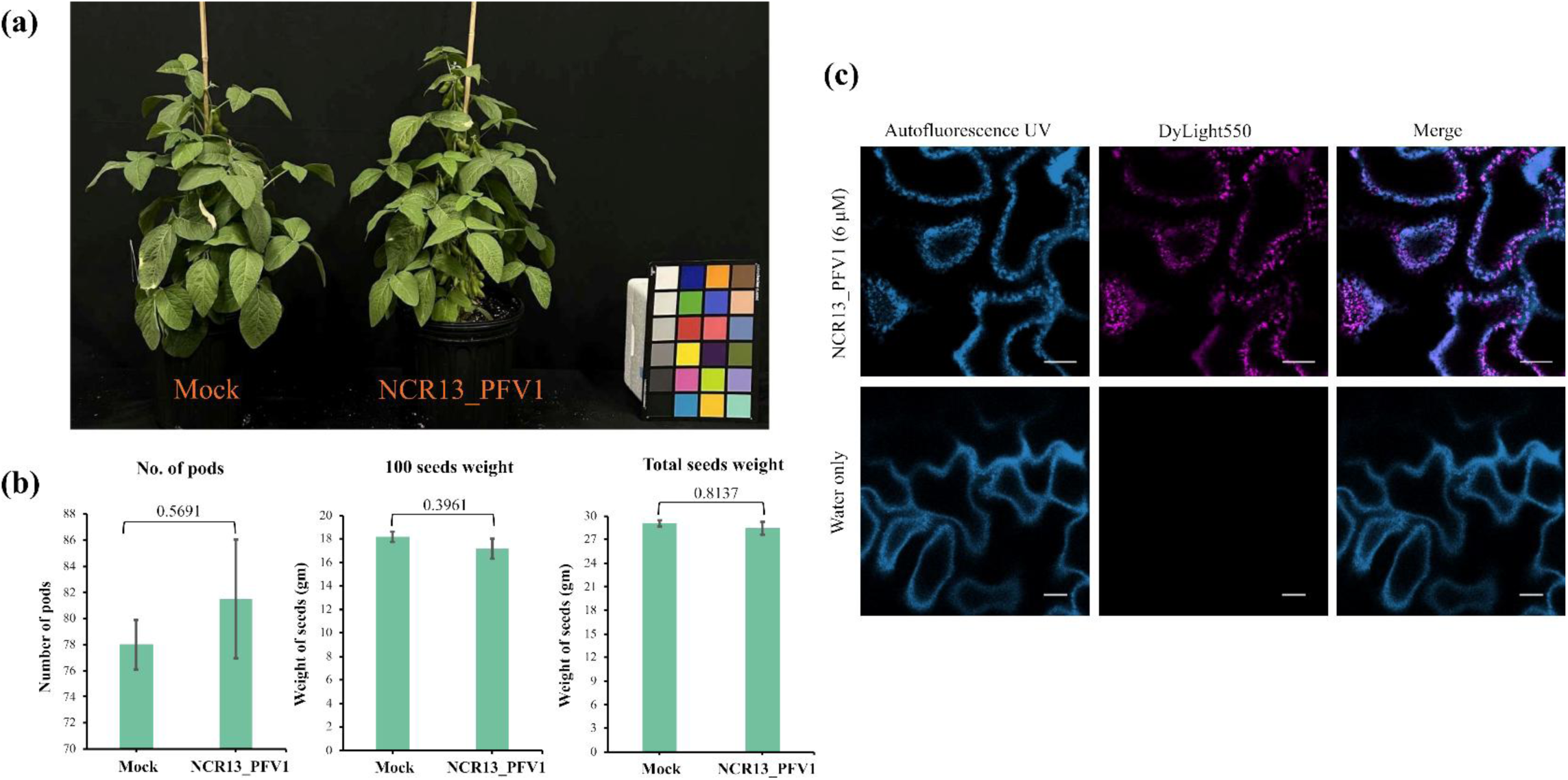
NCR13_PFV1 has no phytotoxic effects on soybean growth and yield. **(a)** Representative picture showing the morphology of 3-month-old soybean plants treated with water (mock) and 6 μM NCR13_PFV1. **(b)** Bar graphs showed the number of pods, 100 seeds weight, and total seed weight of soybean plants treated with water and NCR13_PFV1. Each data point includes the average of at least 4 plants and error bars indicating standard error of the mean between the replicates. The statistical test was performed using the Student’s *t*-test and the *p*-values among the compared groups are indicated. **(c)** Confocal microscopy images of soybean leaf surface sprayed with 6 μM DyLight550-NCR13_PFV1 or water. Soybean leaves were sprayed with 6 μM fluorophore DyLight550-NCR13_PFV1 and imaged after 24 hr and the peptide remained on leaf surface (not internalized). Scale bars = 10 μm.

## Discussion

FLS is a major foliar fungal disease that threatens soybean production in the United States and other soybean growing regions globally. In view of rapidly emerging pathogen resistance to QoI fungicides, effective management of FLS has become a major challenge, exacerbated by a lack of resistant cultivars. Therefore, discovery of new sustainable bio-fungicides with multiple MoA capable of rescuing the activity of QoI fungicides and mitigating fungicide resistance is a high priority.

Small cysteine rich plant AFPs with multi-faceted MoA and low environmental impact offer promising alternatives to chemical fungicides. To our knowledge, such peptides have not been tested for control of FLS in soybean as spray-on fungicides. In this study, we tested several AFPs for their antifungal activity *in vitro* against *C. sojina* and examined the potential of one of these peptides, NCR13_PFV1, for control of FLS. Its multi-site MoA characterized in our study makes it unlikely that *C. sojina* will readily evolve resistance in the field.

Short chain synthetic peptides spanning the γ-core motif of plant defensins MtDef4, MtDef5, MsDef1 and OefDef1 as well as two NCR13 disulfide cross-linking variants were screened for *in vitro* antifungal activity against *C. sojina*. This study is the first to report antifungal activity of these peptides against *C. sojina*. All defensin-derived peptides tested inhibited the pathogen growth with MIC values ranging from 1.5 to 6 µM. However, NCR13_PFV1 was highly potent against this pathogen with the MIC value of 0.187 µM. In contrast, NCR13_PFV2 with an identical amino acid sequence but different disulfide pairing was eight to sixteen-fold less potent. This was not surprising since NCR13_PFV1 was previously reported to be several fold more potent than NCR13_PFV2 against other ascomycete necrotrophic pathogens (Godwin et al., 2024). Interestingly, NCR13_PFV1 showed similar potency against both QoI-sensitive and - resistant isolates of *C. sojina* indicating its MoA is different from that of these fungicides.

Our study demonstrated that NCR13_PFV1 can be used as a bio-fungicide to control FLS and has no deleterious effects on soybean grown and yield under the conditions applied. Spray application of 6 µM NCR13_PFV1 applied pre- or post-inoculation on soybean plants conferred robust control of FLS caused by the QoI-sensitive and -resistant isolates. The observed reduction of fungal biomass in the peptide-treated leaves further confirmed the efficacy of this peptide in controlling FLS. Because of its highly potent broad-spectrum antifungal activity against several fungal pathogens (*B. cinerea, F. virguliforme*, and *S. sclerotiorum*) (Godwin et al., 2024), NCR13_PFV1 has the potential to be developed as an all-in-one bio-fungicide targeting multiple soybean fungal pathogens. Nevertheless, the efficacy of NCR13_PFV1 to control these fungal diseases must be thoroughly evaluated in the field and any potential off target effects must be identified and addressed.

We previously reported multiple MoA of NCR13_PFV1 against *B. cinerea* (Godwin et al., 2024). In this study, we observed striking similarity in NCR13_PFV1 MoA against *C. sojina*. Specifically, this peptide permeabilized the plasma membrane of *C. sojina* conidia and germlings within 10 min of treatment, as reported earlier for *B. cinerea* (Godwin et al., 2024). However, we observed some differences in the subcellular localization pattern of this peptide in *C. sojina* compared to *B. cinerea*. In *C. sojina*, upon entry NCR13_PFV1 immediately showed unidentified high intensity regions in the cytoplasm. It is unclear if this signal required interaction of the peptide with specific cytoplasmic targets. In contrast to *B. cinerea*, we observed subcellular localization of the peptide into nuclei at very low frequency. As reported previously, NCR13_PFV1 bound to *B. cinerea* rRNA and inhibited protein translation *in vitro.* In this study, we confirmed the binding of NCR13_PFV1 to *C. sojina* rRNA *in vitro* and inhibiton of protein translation in the fungicide resistant *C. sojina* Cs350 *in vivo*. Thus, inhibition of protein synthesis may be an important step in the MoA of this peptide. Further research is needed to identify precise binding sites of this peptide in rRNA and/or specific ribosomal proteins.

One of the major challenges in controlling FLS disease in soybeans is the development of resistance to QoI fungicides in *C. sojina* (Zeng et al., 2015). Therefore, it is imperative that current research in FLS disease management focuses on alternative, environmentally friendly sustainable solutions. Combining chemical fungicides with AFPs can reduce the likelihood of fungi developing resistance to “at risk” fungicides. This strategy is particularly promising if peptide-fungicide combination works additively or synergistically to improve the efficacy of fungal diseases management. In our study, checkerboard assays, confocal microscopy, as well as *in planta* antifungal assays revealed that NCR13_PFV1 and QoI fungicides exhibit additive antifungal activity against *C. sojina* to reduce FLS disease severity in soybean. Confocal microscopy studies showed that QoI fungicide azoxystrobin leveraged the early plasma membrane permeabilization ability of NCR13_PFV1 to enter and kill the *C. sojina* cells more effectively and rapidly, even at lower concentrations. As the MoA of QoI fungicides against *C. sojina* involves the inhibition of mitochondrial respiration, it is plausible that the peptide-fungicide combination could achieve more efficient mitochondrial respiration inhibition; however, this hypothesis requires experimental validation. These findings are exciting and have significant potential for the development of peptide-fungicide formulations to control a variety of plant diseases, including FLS in soybeans.

Differential gene expression analysis of NCR13_PFV1 treated *C. sojina* showed that approximately 56.5% of the DEGs were upregulated and 43.5% of DEGs were downregulated in *C. sojina* after 30 min of peptide treatment. Of the top five significantly upregulated genes, one gene (g7198) was predicted to have functions related to oxidative burst. This corresponded with our observation of peptide-induced ROS in *C. sojina*. The GO enrichment analysis identified that NCR13_PFV1 induced the enrichment of GO terms related to heme/iron binding in *C. sojina* and *B. cinerea*. Interestingly, a previous report on a well-studied NCR247 showed that the peptide binds to heme and promotes the uptake of iron in rhizobia (Sankari et al., 2022). Briefly, when NCR247 binds to heme, it creates iron deprivation state, which in turn causes iron starvation response in rhizobia thus resulting in more uptake of iron (Sankari et al., 2022). Our study showed up-regulation of 16 genes encoding heme binding proteins in *C. sojina*. Using the peroxidase assay described by (Sankari et al., 2022), we assessed the heme binding potential of NCR13_PFV1, but found no evidence of heme binding. Even though NCR13_PFV1 does not directly bind to heme (as tested *in vitro*), it may still induce a similar iron starvation response *in vivo* like NCR247 and promote iron uptake in *C. sojina*. *In vitro* antifungal activity studies of NCR13 variants against *C. sojina* and *B. cinerea* at various concentrations of iron revealed that the fungus becomes more tolerant to the peptide with higher concentration of iron. Additionally, increased iron concentration reduced the amount of ROS in *C. sojina*. RNA-seq data revealed the upregulation of genes encoding various heme binding proteins such as catalases, peroxidases, and cytochrome P450s in response to peptide challenge. Catalases and peroxidases require heme to enhance resistance against ROS as shown in various fungi such as *Candida albicans* and *Magnaporthe oryzae* (Pradhan et al., 2017; Zámocký et al., 2020). Cytochrome P450s in fungi are known to be involved in xenobiotic detoxification (Coleman et al., 2011; Shin et al., 2018). Additionally, recent studies in *Fusarium graminearum* showed that deletion of genes encoding heme binding proteins made fungi sensitive to oxidative stress and increased sensitivity to fungicide tebuconazole (Park et al., 2023; Wang et al., 2024). Moreover, heme is also crucial for fungal growth, external stress response, as well as virulence. Overall, these results revealed that NCR13_PFV1 induces ROS and the expression of genes encoding heme binding proteins in *C. sojina*. Iron supplementation resulted in reduced ROS thus alleviating the oxidative stress caused by peptide treatment. This highlights the importance of iron and iron-related genes in fungi for tolerance against peptides. Further investigation is needed to understand how *C. sojina* may develop tolerance to various antifungal peptides including NCR13_PFV1 which is crucial for monitoring potential resistance development in the future.

In summary, the data presented here allowed us to propose the model for the antifungal action of NCR13_PFV1 in *C. sojina* (Fig. 9) and for the control of FLS in soybean. It also highlights the additive antifungal activity between NCR13_PFV1 and QoI fungicides, offering a promising approach for crop protection against FLS through combined foliar application of peptides and QoI fungicides.

**Figure 9:**
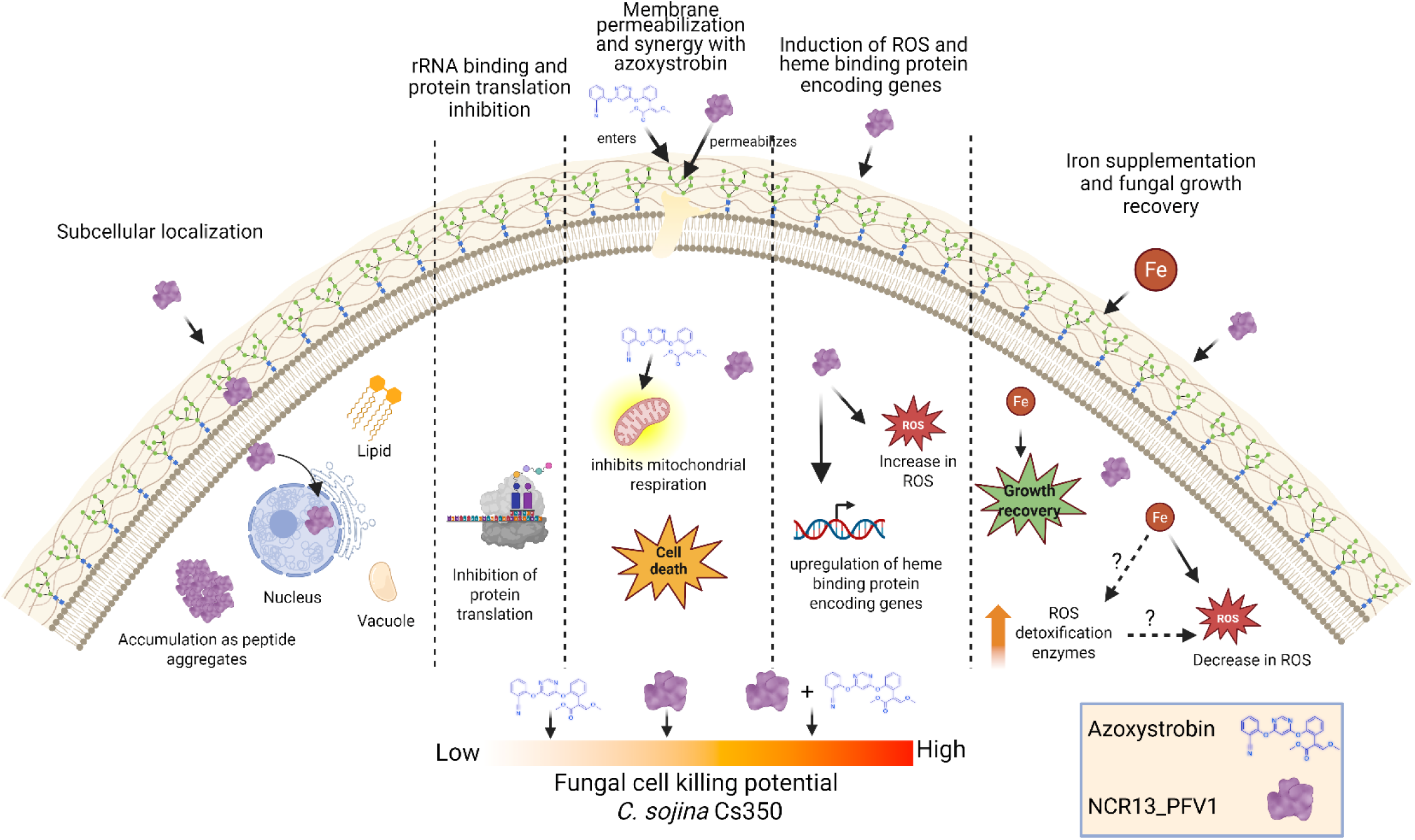
Model illustrating the MoA of NCR13_PFV1 against QoI fungicide resistant *C. sojina* isolate Cs350. The figure depicts the subcellular localization of NCR13_PFV1 in the nucleus and cytoplasm. It shows the peptide’s ability to bind to rRNA and inhibit protein translation. It highlights the membrane permeabilization, additive antifungal activity of NCR13_PFV1 with QoI fungicide azoxystrobin, and overall fungal cell killing potential of NCR13_PFV1, azoxystrobin and their combination. It further demonstrates the induction of ROS and heme binding protein encoding genes in *C. sojina* upon peptide treatment as well as the fungal growth recovery and reduction of ROS upon iron supplementation. Image created using https://www.biorender.com/.

## Materials and Methods

### QoI fungicide sensitive and -resistant *C. sojina* isolates and their growth

*C. sojina* QoI fungicide sensitive (Cs320, Cs324, and Cs752) and -resistant (Cs350, Cs326, and Cs673) isolates obtained from Dr. Carl Bradley, University of Kentucky, were used in this study. These isolates were grown and maintained in 10% clarified V8 juice agar medium (Uppala et al., 2019) at room temperature under 12 h light-12 h dark conditions. The conidia from the fungal isolates were collected in low salt synthetic fungal medium (SFM) (Liang et al., 2001) and adjusted to the required density using hemocytometer.

### Peptide synthesis and purification

AFPs used in this study are listed in Table S6. Expression of NCR13 in *P. pastoris* X33 and its purification were performed as described (Godwin et al., 2024). Chemically synthesized defensin-derived and NCR13-derived short peptides used in this study were obtained from WatsonBio Sciences (Houston, TX) and further purified by HPLC (Sankari et al., 2022; Tetorya et al., 2023; Godwin et al., 2024; Kalunke et al., 2025). The concentration of each purified peptide was determined using the Pierce BCA Protein Assay Kit (Thermo Scientific).

### *In-vitro* antifungal activity assays

*In-vitro* antifungal activity of AFPs and QoI fungicides was determined using 96-well plate assays as described (Velivelli et al., 2020) with slight modifications. Briefly, 2 to 3-week-old *C. sojina* was used to make a conidial suspension in 2X SFM media at the final concentration of ∼10^5^ conidia mL^-1^. Simultaneously, peptide dilutions (in water) at various concentrations were made. Fungicides were dissolved in 50% acetone and 50% water to make a stock solution of 500 µM and serial dilutions were made in water as needed. The peptide-conidia or the fungicide-conidia mixture at the ratio of 1:1 (45 µl:45 µl) was added in a 96-well plate and incubated at room temperature for 48 hr after which MICs of the peptides/fungicides were calculated based on the microscopy as well as resazurin cell viability assay (Velivelli et al., 2020).

### *In planta* antifungal assays and fungal DNA quantification using qPCR

For both pre- and post-inoculation *in planta* antifungal activity assays, 2 to 3-week-old soybean plants cv. Blackhawk (FLS susceptible) grown under 16 hr light and 8 hr dark cycle were used. For pre-inoculation *in planta* antifungal activity assay, the second or third trifoliate leaves of soybean plants were first sprayed with 1 mL of NCR13_PFV1 peptide (6 µM) in the abaxial leaf surface followed by spray inoculation with 1 mL of conidial suspension (1 x 10^5^ mL^-1^) of Azoxystrobin-sensitive Cs320 or -resistant Cs350 isolate. For the post-inoculation *in planta* antifungal activity assay, the second or third trifoliate leaves of soybean were first spray-inoculated with 1 mL of conidial suspension (1 x 10^5^ mL^-1^) followed by spray application of 1 mL of NCR13_PFV1 peptide (6 µM) 2 hr later. The mock plants were sprayed with water but no peptide. For both pre- and post-inoculation *in planta* assays, each treatment consisted of at least 4 plants and the experiments were performed at least two times. The inoculated plants were kept in a growth chamber under 16 hr light and 8 hr dark cycle and 90% relative humidity. The FLS symptoms started to appear 7-10 days post-inoculation and disease scoring was done at 14 days. Diseased leaves were excised from the plant and both white light and CropReporter (Fv/Fm) images were taken to assess the photosynthetic efficiency of the plant. The leaves were also scored based on 0-5 scale where: 0 – healthy leaves with no leaf spots, 1 – 1%-25% leaf area covered with spots, 2 – 26%-50% leaf area covered with spots, 3 – slight yellow leaves with 51%-75% leaf area covered with spots, 4 – moderate yellow leaves with 76%-95% leaf area covered with spots, 5 – whole leaf is yellow or >96% leaf area covered with spots.

Furthermore, DNA was extracted from the excised leaves to quantify the fungal DNA content using qPCR. Two genes *con6* (CD397253) and *con7* (AW310136) (were used as reference control genes for soybean (Libault et al., 2008) and ITS (internal transcribed spacer, MZ451112), Histone H3 gene (MZ456946) were used as a housekeeping control gene for *C. sojina*. DNA quantification in terms of fold change was calculated using ΔΔCt method (Livak and Schmittgen, 2001). The primers used for qPCR to amplify soybean and *C. sojina* control genes are listed in Table S7. Significant differences in the disease severity values and the fungal DNA content between the treatments were determined using the Wilcoxon signed-rank test in *R*. The compared treatments were considered significantly different at *p*-value ≤0.05.

### Peptide-fungicide synergy using checkerboard assays

Synergy between NCR13_PFV1 and widely used QoI foliar fungicides, azoxystrobin (Sigma-Aldrich, Cat No:31697), trifloxystrobin (Sigma-Aldrich, Cat No:46447), and pyraclostrobin (Sigma-Aldrich, Cat No:33696) were evaluated using checkerboard assays. This assay was performed in a 96-well plate and the preparation of conidial suspension, peptide dilutions, and fungicide dilutions was similar to the *in-vitro* antifungal activity assay described above. The presence of alternative respiration pathway allows some fungi including *C. sojina* to germinate even in the presence of high dose of QoI fungicides *in vitro* (Ziogas et al., 1997; Zhang et al., 2012). Therefore, to prevent alternative respiration in *C. sojina*, salicylhydroxamic acid (SHAM) was added in each reaction at the final concentration of 60 µg/mL (Zhang et al., 2012). The conidia, peptide, fungicide, and SHAM were added in a 96-well plate at the v/v ratio of 4:1:1:2 (50 µl: 12.5 µl: 12.5 µl: 25 µl) in the final reaction volume of 100 µl and the plate was incubated at room temperature for 96 hr (4 days). After 96 hr, MIC values of the peptide-fungicide combination, peptide alone and fungicide alone were calculated based on the microscopy as well as the resazurin cell viability assay. The fractional inhibitory concentration (FIC) index was calculated using the formula FIC = (MICA+B/MICA) + (MICB+A/MICB), where MICA and MICB is the MIC for peptide (A) alone and fungicide (B) alone respectively, MICA+B is the MIC for peptide in combination with fungicide (A+B), and MICB+A is the MIC for fungicide in combination with peptide (B+A). The FIC index was interpreted as Synergistic: ≤ 0.5; Additive: > 0.5-1.0; Indifferent: >1-< 4.0; Antagonistic: ≥ 4.0.

### Cell death detection assay

The fungicidal activity of azoxystrobin and peptide NCR13_PFV1 in an azoxystrobin-resistant *C. sojina* Cs350, was analyzed by visualizing the influx of a cell death indicator dye propidium iodide (PI) using timelapse confocal microscopy. The *C. sojina* conidia were treated with various concentrations of azoxystrobin, or NCR13_PFV1, or azoxystrobin plus NCR13_PFV1, or water and 0.5 μg/mL of PI, and were observed under confocal microscopy at an excitation and emission wavelength of 500 nm and 545-690 nm respectively. Z-stacks at each timepoint were acquired using 63X water immersion objective lens (HC PL Apochromat CS2). For each sample, the number of live and dead cells after one hour of peptide or fungicide or combination treatment was determined.

### Plasma membrane permeabilization assay

Membrane permeabilization of *C. sojina* Cs350 by NCR13_PFV1 and azoxystrobin was analyzed by visualizing the influx of SYTOX Green (SG) (Thermo Scientific) using time-lapse confocal microscopy. Germlings grown 6-8 hr or conidia of *C. sojina* were mixed with various concentrations of NCR13_PFV1 or azoxystrobin or both and 0.5 µM SG and were observed under confocal microscopy at an excitation wavelength of 500 nm and an emission wavelength ranging from 510 to 600 nm at specific time intervals. A z-stack at each timepoint was acquired using 63X water immersion objective lens (HC PL Apochromat CS2).

### *In planta* antifungal assays to determine synergy between NCR13_PFV1 and the QoI fungicide azoxystrobin

To further test additive/synergistic interaction between NCR13_PFV1 and azoxystrobin against the fungicide resistant isolate Cs350, *in planta* assays were performed. These assays were similar to the post-inoculation *in planta* assays as mentioned above. Briefly, second or third trifoliate leaves of soybean were first sprayed with 1 mL of Cs350 conidial suspension (1 x 10^5^ mL^-1^) followed by spray application with 1 mL of NCR13_PFV1 (3 µM) or azoxystrobin (3 µM or 6 µM) or their combination (3 µM NCR13_PFV1 plus 3 µM azoxystrobin or 3 µM NCR13_PFV1 plus 6 µM azoxystrobin) or water after 2 hr of pathogen inoculation. Plants were incubated in the growth chamber and disease scoring was done as described above.

### Uptake of NCR13_PFV1 by *C. sojina*

NCR13_PFV1 was labeled with DyLight550 amine-reactive dye following the manufacturer’s instructions (Thermo Scientific) and time-lapse confocal microscopy was performed to measure the uptake and subcellular localization of the peptide in *C. sojina* Cs350 cells. The nuclear staining dye Hoechst 3352 was used along with DyLight550-NCR13_PFV1 to determine the subcellular localization of NCR13_PFV1 in *C. sojina*. Conidia grown for 6-8 hr were treated with 0.09 µM of Dylight550-NCR13_PFV1 and incubated for 30 min after which 4 µM Hoechst 3352 dye was added for another 15 min. Treated conidia were used to capture z-stacks at specific time intervals using a Leica SP8-X confocal microscope (63X water immersion objective lens, HC PL Apochromat CS2) in fluorescence and transmitted light mode. The excitation and emission wavelengths of DyLight550-NCR13_PFV1 and Hoechst 3352 were 550, 560-660, and 405, 420-500 nm respectively. Similarly, to determine the subcellular localization of NCR13_PFV1 in other organelles, *C. sojina* conidia were also stained with vacuole-specific dye CMAC and lipid staining dye BODIPY 493/503 (Thermo Scientific). The excitation and emission wavelengths for CMAC and BODIPY were 405, 420-500, and 493, 505-540 nm respectively. Images were displayed as single optical section or maximum intensity projections.

### Electrophoretic mobility shift assay (EMSA)

To determine the RNA binding ability of NCR13_PFV1, EMSA was performed as described in (Godwin et al., 2024). Briefly, total RNA was extracted from *C. sojina* Cs350 grown in SFM media using ZymoBIOMICS DNA/RNA Miniprep Kit (Zymo Research). Approximately 200 ng of the fungal RNA was mixed with different concentrations of NCR13_PFV1 in 20 μl of binding buffer (10 mM Tris-HCl, pH 8.0, 1 mM EDTA), incubated at room temperature for 1 hr, and ran through gel electrophoresis.

### *In vivo* protein translation inhibition assay

To assess the translation inhibition of NCR13_PFV1 in *C. sojina* Cs350, we adapted a recently established puromycin labelling protocol in *B. cinerea* with minor modifications (Godwin and Shah, 2025). *C. sojina* spores at a concentration of approximately 10⁵/mL were inoculated in SFM medium and cultured for 18 hr. The germlings were then treated with NCR13_PFV1 (0.75 and 1.5 µM) or cycloheximide (positive control) for 2 hr, followed by the addition of puromycin (1 mM) and incubation for 3 hr at 28°C. After incubation, the germlings were washed three times with sterile water. Total proteins were extracted from germlings using lysis buffer (100mM HEPES, 5 mM EDTA, 0.1% Triton-X 100 and yeast/fungal protease inhibitor cocktail) and the concentration was estimated using Pierce BCA Protein Assay Kit (Thermo Scientific). Equal amounts (10 µg) of fungal protein were separated on a 4%–20% mini-PROTEAN TGX stain-free protein gel (Bio-Rad, Cat No: 4568094). Western blot analysis was performed using an anti-puromycin antibody (Cell Signaling Technology, Cat No: 40939S), and signal intensities were quantified using Fiji (Schindelin et al., 2012).

### RNA extraction, sequencing, and transcriptomic analysis

Conidia from two-week-old *B. cinerea* and fungicide resistant *C. sojina* Cs350 were transferred to 50 mL Erlenmeyer flask with 5 mL of 1X SFM media at the final concentration of approximately 10^5^ conidia. The conidia were then grown in shaker in 200 rpm at 28°C for 48 hr after which 2.5 mL mycelial culture was collected and flash frozen in liquid nitrogen and stored at -80°C (Control, timepoint 0 min/ T0). To the remaining 2.5 mL mycelial suspension, NCR13_PFV1 or NCR13_PFV2 was added at the final concentration of 0.187 µM for *C. sojina* and 0.09 µM for *B. cinerea*. After 30 min of peptide challenge, the mycelial sample was flash-frozen in liquid nitrogen and stored at -80°C (Treated, timepoint 30 min/ T30). RNA extraction was done using ZymoBIOMICS DNA/RNA Miniprep Kit (Zymo Research). cDNA library preparation, mRNA sequencing, and data filtering to obtain clean reads were performed at Novogene (California, USA).

The gene prediction in *C. sojina* genome (Gu et al., 2020) was performed using Augustus 3.4.0 (Stanke et al., 2006). As for *B. cinerea* RNA-seq data, the reference genome and annotation files of *B. cinerea* T4 were used for analysis (Staats and van Kan, 2012). The raw RNA-seq reads were cleaned by removing low-quality reads, N>10% and reads containing adapters. Clean RNA-seq reads were mapped to the genome using HISAT2 (Kim et al., 2019) and the aligned reads were assembled using StringTie assembler (Pertea et al., 2015). The gene expression levels and number of transcripts mapped to the genome were determined using featureCounts (Liao et al., 2014). The data normalization and differential gene expression analysis was done using DESeq2 (Love et al., 2014). The gene enrichment analysis, GO and KEGG enrichment were done using the clusterProfiler (Yu et al., 2012). GO terms and KEGG pathways with padj < 0.05 were considered significantly enriched. Gene Set Enrichment Analysis (GSEA) (Subramanian et al., 2005) was done to determine differences in gene sets between no peptide and peptide treatments.

### Heme preparation, heme-peptide binding assay, and peroxidase assay

Heme solutions were freshly prepared by dissolving hemin chloride (Sigma Cat No: 3741) in 0.1M NaOH and used within an hour after preparation. Two heme sequestering peptides, NCR247 and NCR455 (Sankari et al., 2022) were used as positive controls, and water only treatment as a negative control. The equimolar concentration of heme and peptide (100 µM) or water were mixed and incubated in room temperature for 1 hr. When dissolved in NaOH heme is light to dark brown in color, however when it binds to a peptide the color changes to reddish-brown as described by (Sankari et al., 2022). The change in color of heme to reddish-brown indicated the heme binding ability of the peptide.

A Pierce TMB substrate kit (Thermo Scientific) was used to measure the peroxidase activity of heme. In a transparent 96-well plate, 50 µl of TMB substrate and 50 µl of peroxidase solution were added followed by addition of heme (5 µM) or peptides plus heme (5 µM each). The reaction progression was measured in the Tecan plate reader, by recording UV-vis absorption values at 370 nm and 652 nm for every minute until one hour.

### Measurement of ROS in *C. sojina* upon NCR13_PFV1 treatment

ROS indicator dye 2′,7′-dichlorodihydrofluorescein diacetate (H2DCFDA) was used to evaluate ROS induction in *C. sojina* cells. *C. sojina* conidia (∼1 x 10^5^ conidia mL^-1^) in SFM media at various concentrations of iron (FeSO4) was mixed with 15 µM H2DCFDA, 0.18 µM NCR13_PFV1, and the fluorescence measurements were taken in a Tecan plate reader M200 PRO at excitation and emission wavelengths of 485 and 535 nm respectively at every 30 min for 15 hr. ROS induction in *C. sojina* was visualized using time lapse confocal microscopy (63X water immersion objective lens, HC PL Apochromat CS2). *C. sojina* conidia grown for 4-5 hr in SFM media at various concentrations of iron were treated with 15 µM H2DCFDA and 0.187 µM DyLight550-NCR13_PFV1 followed by acquisition of z-stacks using Leica SP8-X confocal microscope. The excitation and emission wavelengths of DyLight550-NCR13_PFV1 and H2DCFDA were 560, 575-650, and 485, 500-540 nm, respectively.

### Assessment of the effects of NCR13_PFV1 spray on the growth and yield of soybean

To test the potential phytotoxicity of NCR13_PFV1 spray on soybean plants, 6 µM NCR13_PFV1 or water was first sprayed on 4-week-old soybean cv. Blackhawk plants (2 mL/plant) followed by biweekly sprays at 5 mL/plant until plants were 10 weeks old. When plants reached 12 weeks, photographs were taken for phenotypic analysis and allowed to mature.

Soybean pods were harvested when the plants were all dry and mature and the number of pods, 100 seed weight, and total seed weight were determined. At least four plants per treatment were tested.

### Distribution of NCR13_PFV1 on the soybean leaf surface

The distribution of DyLight550-NCR13_PFV1 on the soybean leaf surface was analyzed by spraying soybean leaves with 6 µM DyLight550-NCR13_PFV1 and imaging on Leica SP8-X confocal microscope (63X water immersion objective lens, HC PL Apochromat CS2) after 24 hr. The excitation and emission wavelengths for DyLight550 were 550 nm and 560-620 nm and for autofluorescence of soybean cells were 405 nm and 420-470 nm respectively.

## Data availability

The RNA-seq data from *Cercospora sojina* and *Botrytis cinerea* are available in the Sequence Read Archive (SRA) repository of National Center for Biotechnology Information (NCBI) with accession number PRJNA1247002.

## Author Contributions

Conceptualization, D.M.S and A.P; Methodology, A.P, V.S.N, J.G, R.M.K and M.T.; Formal analysis and investigation, A.P; Writing—original draft, A.P; Writing—review & editing, A.P, J.G, V.S.N, C.J.K. and D.M.S; Supervision, D.M.S and C.J.K.; Funding acquisition, D.M.S.

## Acknowledgements

We are grateful to Dr. Carl Bradley of the University of Kentucky for providing the QoI fungicide sensitive and resistant isolates of *C. sojina*. We thank DDPSC Plant Growth Facility for their help in growing and maintaining plants, and Plant Phenotyping Facility for their help in phenotyping diseased soybean leaves. We also thank Dr. Arnaud Thierry Djami-Tchatchou for his initial help with plant work. We also acknowledge the Advanced Bioimaging Laboratory (RRID: SCR_018951) at the Donald Danforth Plant Science Center (DDPSC) for imaging support, and the use of the Leica SP8-X confocal microscope, funded by an NSF Major Research Instrumentation grant (DBI-1337680) and the ZEISS Elyra 7 Super-Resolution Microscope funded by an NSF Major Research Instrumentation grant (DBI-2018962).

