## Supplementary Figures for "NCR13 peptide protects soybean against *Cercospora sojina* by multiple modes of action and additive interaction with chemical fungicides"

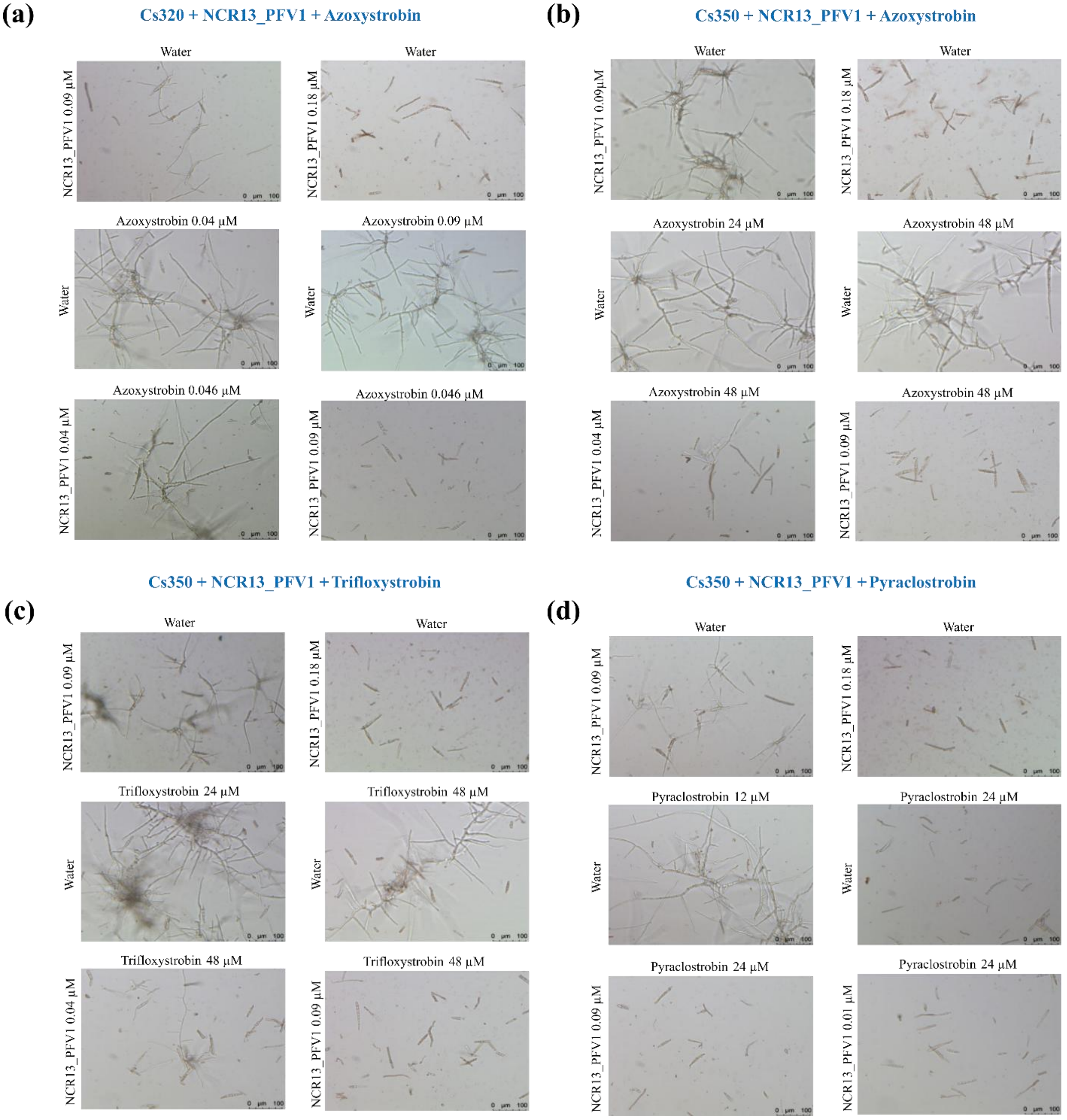


**Fig. S1: NCR13_PFV1 exhibits additive antifungal activity with quinone outside inhibitor (QoI) fungicides against *C. sojina*. (a)** The light microscopy images of fungicide sensitive *C. sojina* isolate Cs320 in various combination of NCR13_PFV1 and Azoxystrobin. Azoxystrobin was tested in the range of 0.01 to 0.09 µM and NCR13_PFV1 was tested in the range of 0.005 to 0.09 µM. The images at MIC and below MIC are shown if applicable. **(b) – (d)** The light microscopy images of fungicide resistant *C. sojina* isolate Cs350 in various combination of NCR13_PFV1 with Azoxystrobin, Trifloxystrobin, and Pyraclostrobin respectively. Fungicides were tested in the range of 1.5 to 48 µM and NCR13_PFV1 was tested in the range of 0.01 to 0.18 µM. The images at MIC and below MIC are shown if applicable.

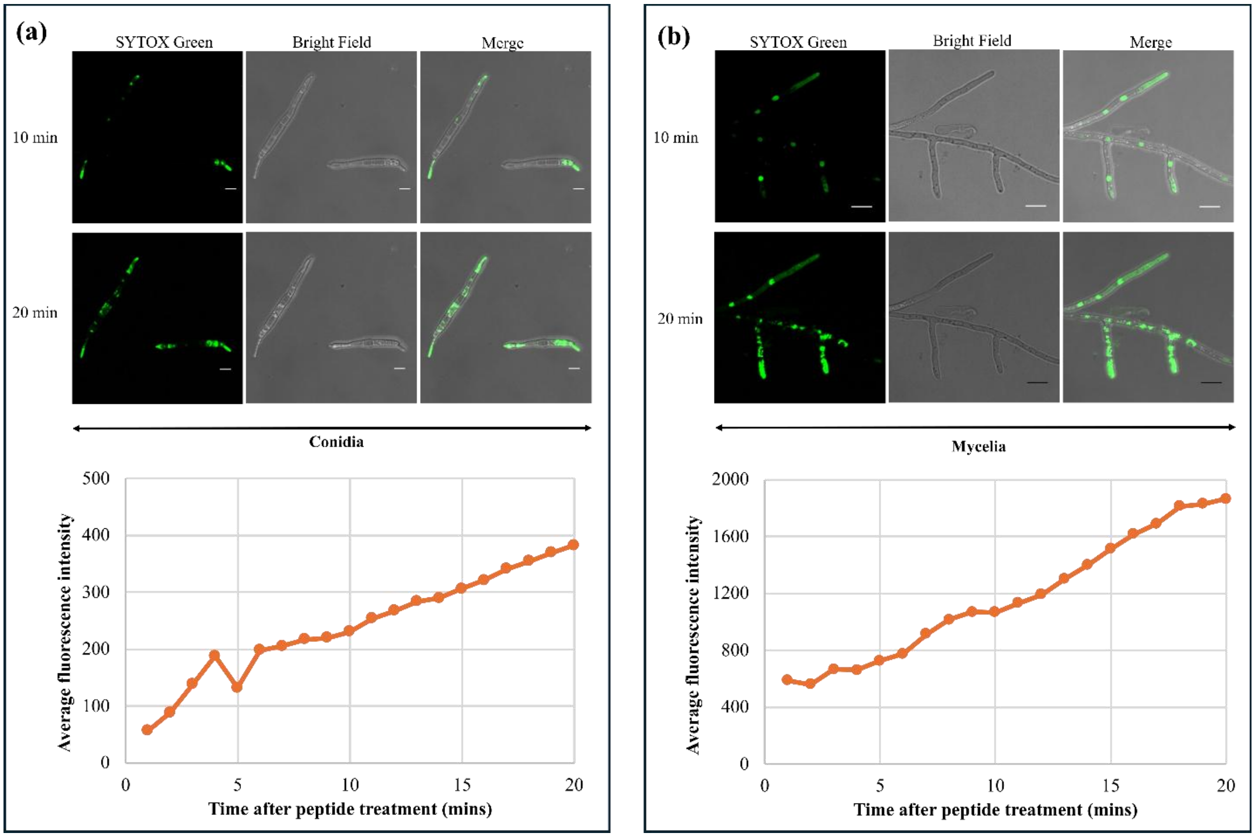


**Fig. S2: NCR13_PFV1 permeabilizes the plasma membrane in *C. sojina*.** Confocal microscopy images of SYTOX Green (SG) uptake in *C. sojina* and their average fluorescence intensity in **(a)** conidia and **(b)** mycelia treated with 0.187 µM of NCR13_PFV1. Time lapse confocal microscopy was performed in *C. sojina* and representative images at 10 and 20 minutes after peptide treatment are shown. Scale bars = 10 µm.

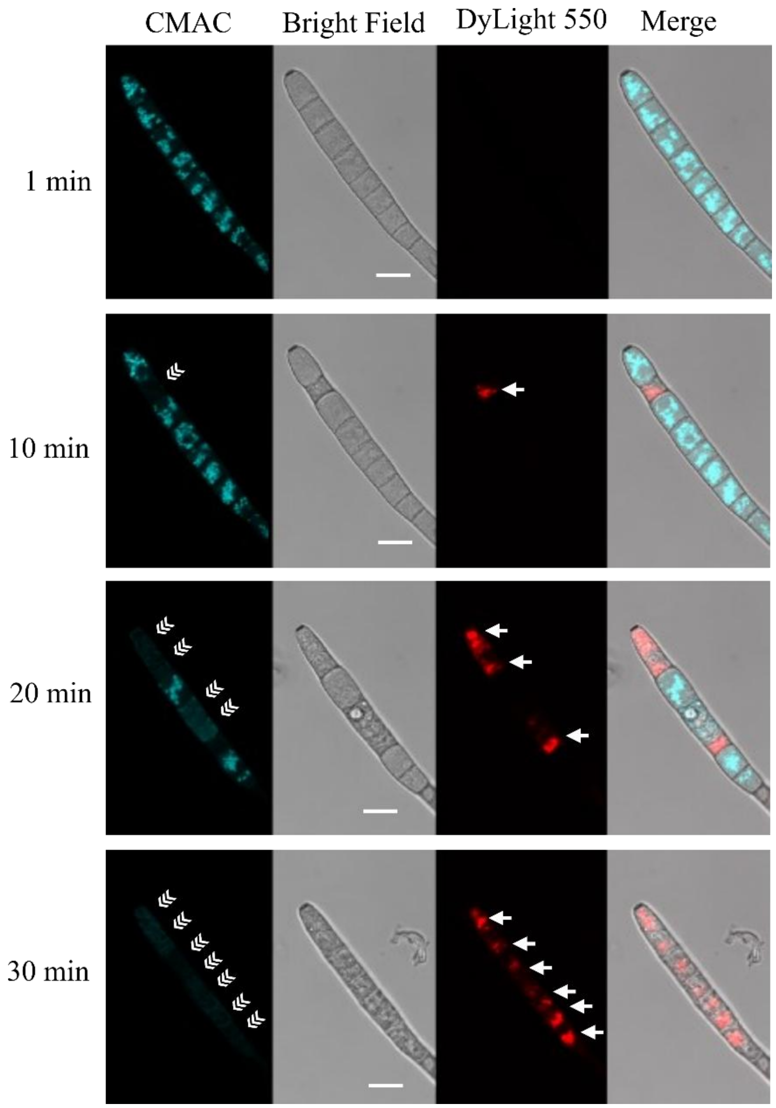


**Fig. S3: NCR13_PFV1 does not localize in vacuoles in *C. sojina* Cs350.** Timelapse confocal microscopy images of *C. sojina* showing the internalization of 0.187 µM DyLight550 labelled NCR13_PFV1. Vacuole specific dye CMAC was used to stain *C. sojina* vacuoles. Triple white arrows indicate the bursting of vacuole, and single white arrows indicate the aggregation of peptide. Scale bars = 10 µm.

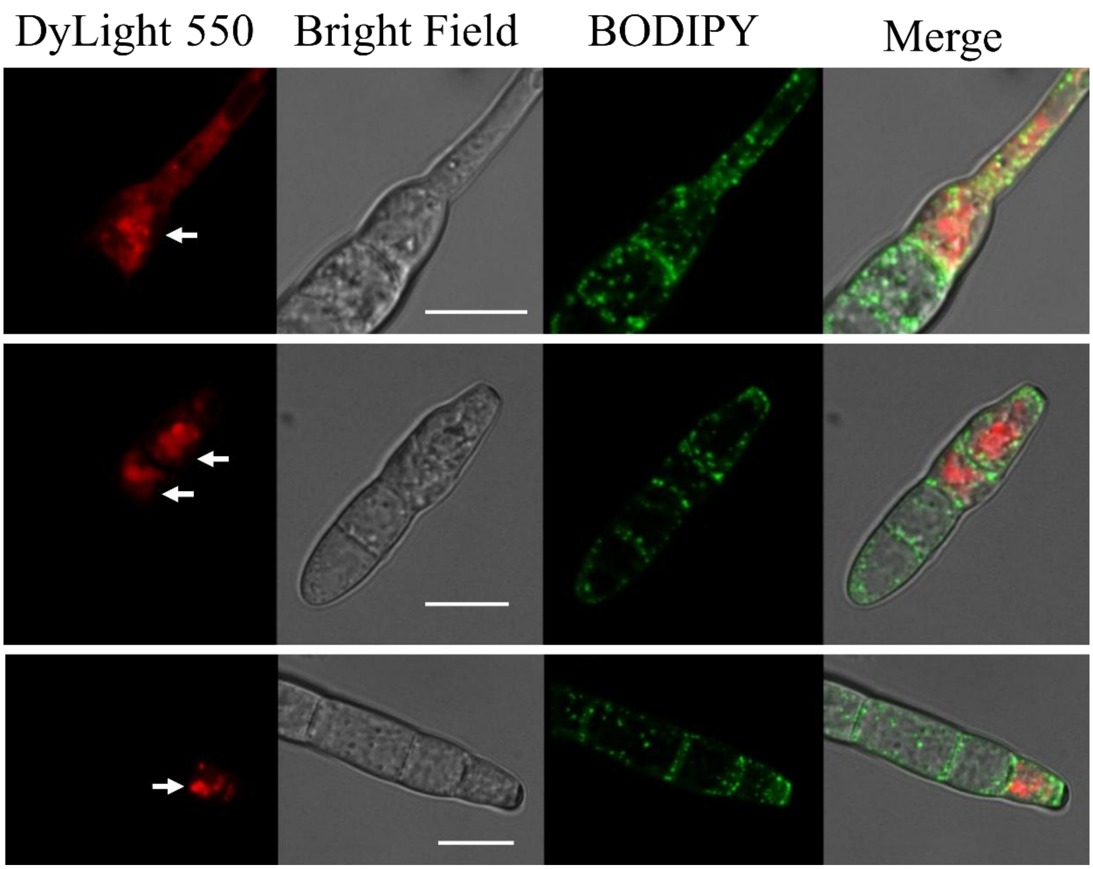


**Fig. S4: NCR13_PFV1 does not localize in lipids in *C. sojina* Cs350.** Timelapse confocal microscopy images of *C. sojina* showing the internalization of 0.187 µM DyLight550 labelled NCR13_PFV1. Lipid specific dye BODIPY was used to stain *C. sojina* lipids. Single white arrows indicate the aggregation of peptide. Scale bars = 10 µm.

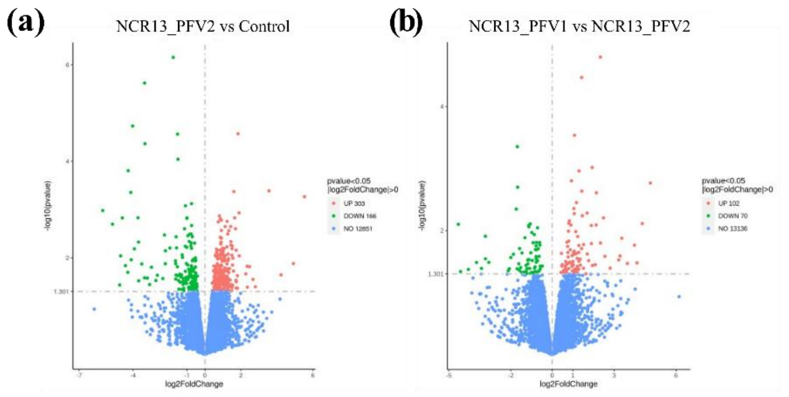


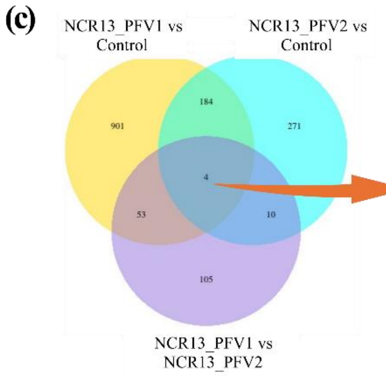


| Gene id | Gene description |
|---|---|
| g12795 | Alpha-dioxygenase 1 \| PF03098:Animal haem peroxidase |
| g7028 | PF14223:gag-polypeptide of LTR copia-type |
| g3617 | Carboxylic acid transporter protein homolog\| PF00083:Sugar (and other) transporter |
| g8753 | - |

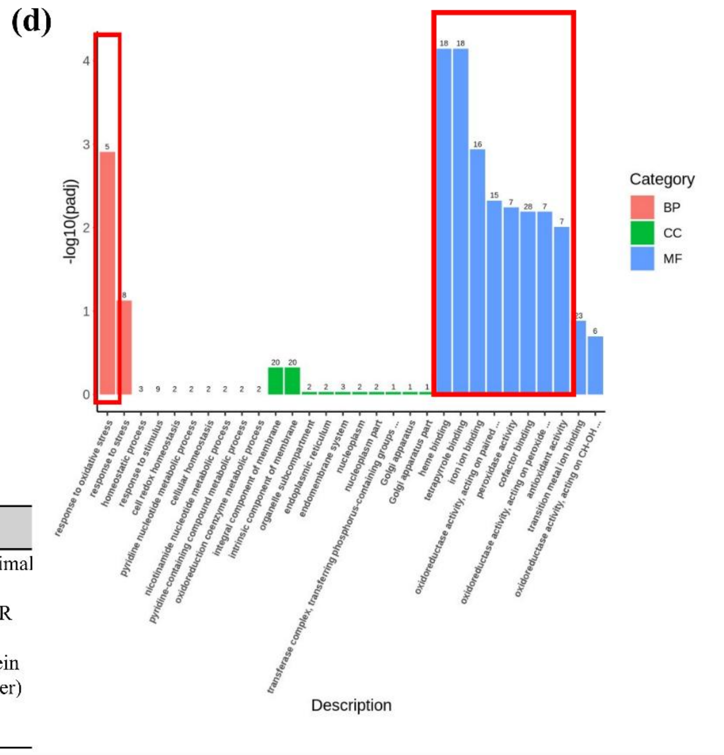


**Fig. S5: NCR13_PFV2 induces genes encoding heme binding proteins in *C. sojina* Cs350.** Volcano plot showing the number of differentially expressed genes (DEGs) in *C. sojina* after 30 min of **(a)** NCR13_PFV2 treatment when compared to the no peptide control at 0 min, and in **(b)** NCR13_PFV1 treatment when compared to NCR13_PFV2 treatment at 30 min. **(c)** Venn diagram showing the number of DEGs at three compared groups. The table shows the list of four DEGs that are common in all three compared groups. **(d)** Bar graph showing the top ten GO terms enriched in each category (BP-Biological Process, CC-Cellular Component, and MF-Molecular Function) at 30 mins after NCR13_PFV2 treatment in *C. sojina*. The significantly enriched GO terms at *p*-value≤0.05 are highlighted in red rectangular box.

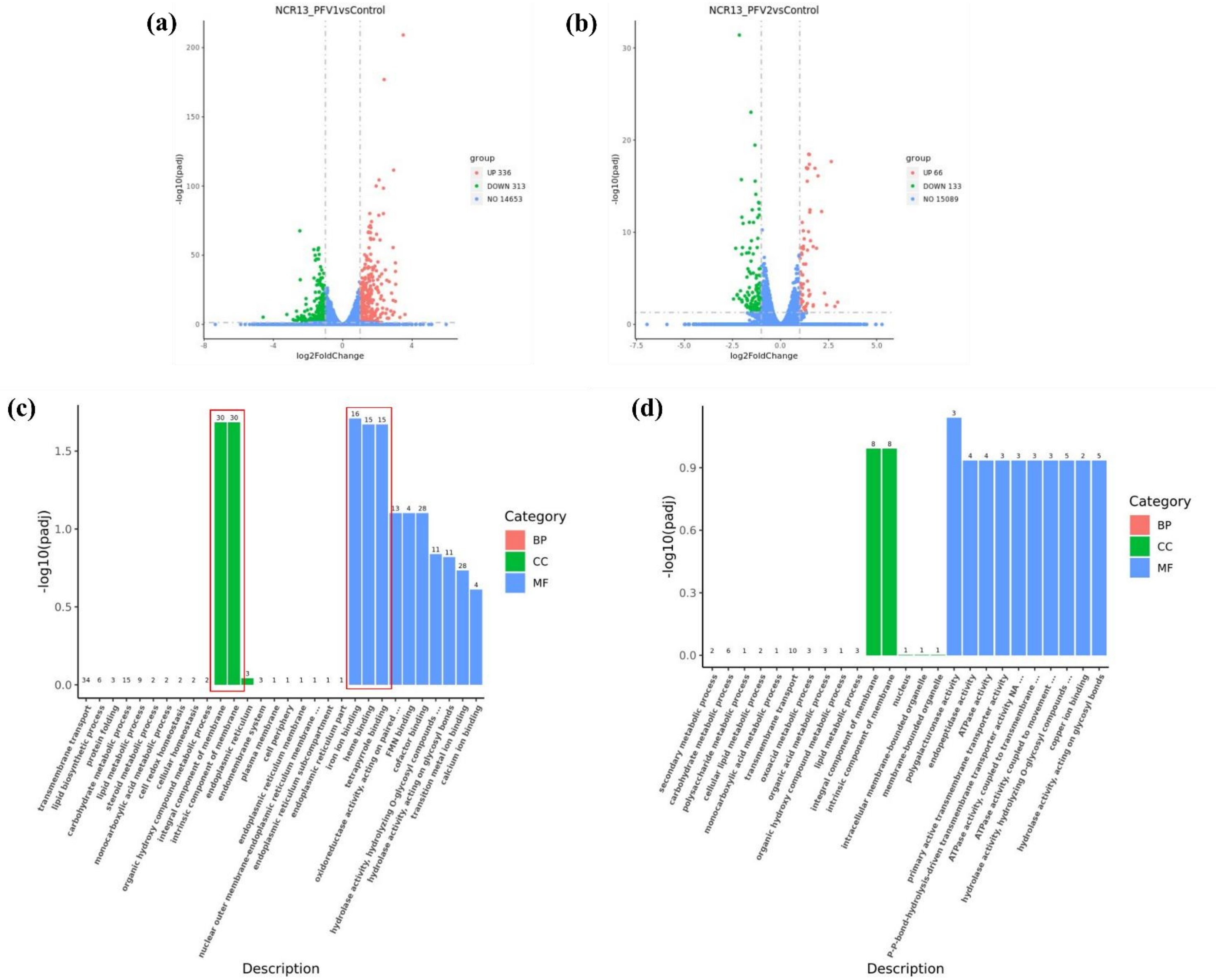


**Fig. S6: NCR13_PFV1 induces genes encoding heme binding proteins in *B cinerea*.** Volcano plot showing the number of differentially expressed genes (DEGs) in *B. cinerea* after 30 min of **(a)** NCR13_PFV1 and **(b)** NCR13_PFV2 treatment when compared to the no peptide control at 0 min. Bar graphs showing the top ten GO terms enriched in each category (BP-Biological Process, CC-Cellular Component, and MF-Molecular Function) at 30 mins after **(c)** NCR13_PFV1 and **(d)** NCR13_PFV2 treatment in *B. cinerea*. The significantly enriched GO terms at $p$-value≤0.05 are highlighted in red rectangular box.

**(a)**

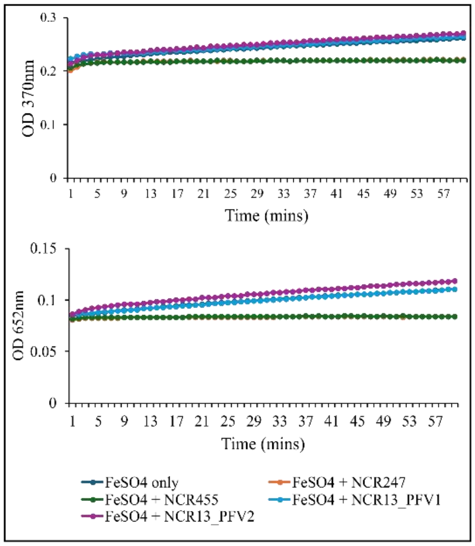


**(b)**

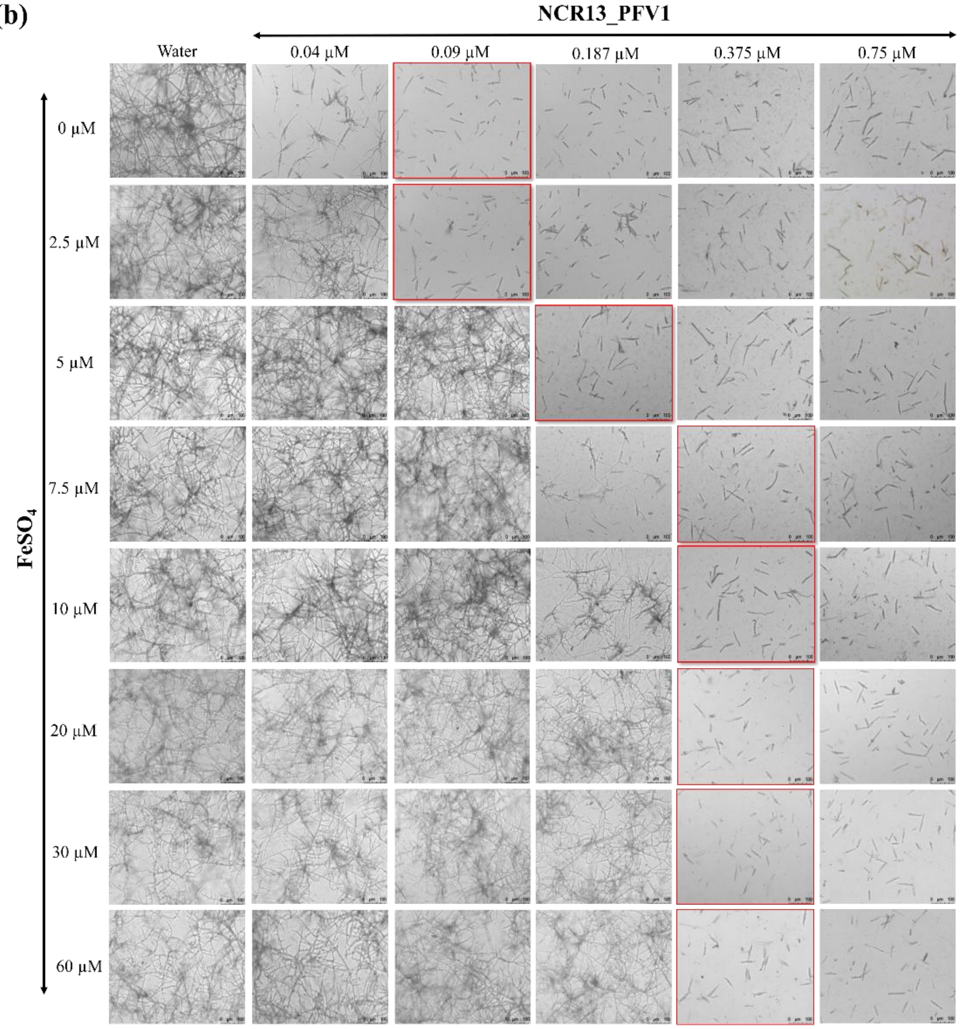


**(c)**

| Iron (FeSO4) concentration (μM) | MIC against *C. sojina* Cs350 (μM) | | |
|---|---|---|---|
| | NCR13_PFV2 | NCR13_V7 | NCR13_V9 |
| **5 (Complete SFM)** | **1.5** | **6** | **3** |
| 20 | 3 | 12 | 6 |

**(d)**

| | Zinc ($ZnSO_4$) concentration (μM) | MIC of NCR13_PFV1 against *C. sojina* Cs350 (μM) |
|---|---|---|
| **Complete SFM** | **0.5** | **0.187** |
| | 2.5 | 0.187 |
| increased | 5 | 0.187 |
| | 10 | 0.187 |

**Fig. S7: *C. sojina* Cs350 exhibits iron mediated tolerance to NCR13 and its variants. (a)** Peroxidase activity of $FeSO_4$ (75 μM) and peptides, NCR247, NCR455, NCR13_PFV1, and NCR13_PFV2 on a chromogenic substrate, 3,3′,5,5′-tetramethylbenzidine (TMB), measured by absorption at 370 nm and 652 nm. **(b)** Light microscopy images of *C. sojina* at various concentration of NCR13_PFV1 and $FeSO_4$. The images highlighted in red indicate the MIC level of *C. sojina* at the indicated concentration of NCR13_PFV1 and $FeSO_4$. **(c)** MIC of NCR13 variants against *C. sojina* at various concentration of iron ($FeSO_4$). **(d)** MIC of NCR13_PFV1 against *C. sojina* at various concentration of zinc ($ZnSO_4$).

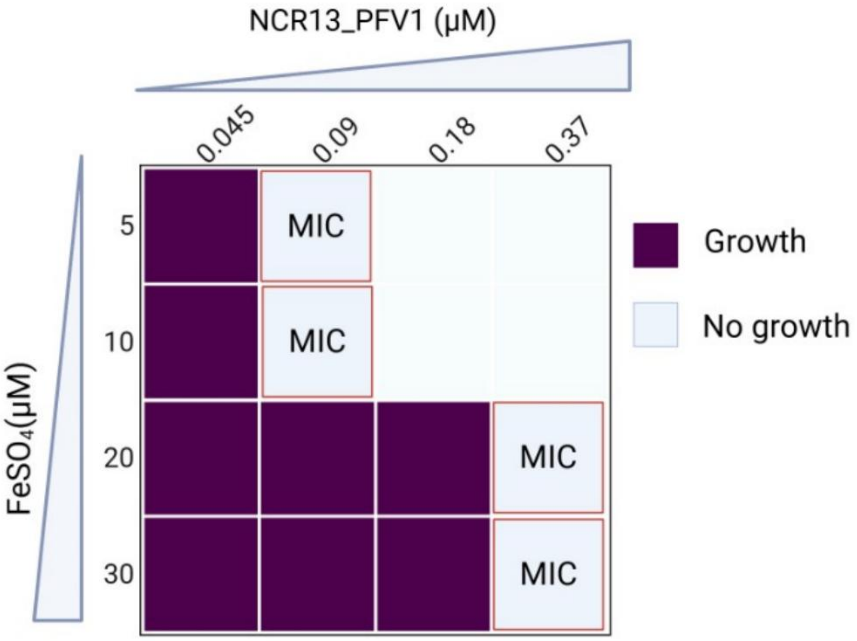


**Fig. S8: *B. cinerea* exhibits iron mediated tolerance to NCR13_PFV1.** MIC of NCR13_PFV1 against *B. cinerea* at various concentration of iron ($FeSO_4$).
